# Spatially resolved transcriptional programs link fallopian tube precursor lesions to immune activation and stromal reorganization

**DOI:** 10.64898/2026.08.27.747392

**Authors:** Maria Sol Recouvreux, Kathleen Trang, Alexandra H. Smick, Shinya Kusumoto, Daniel Pintard, Yael Raz, Barbie-Taylor Harding, Jun Han Kim, Ryan Mika, Michael T. Richardson, Rachel Furge, Dmitrijs Lvovs, Elana Fertig, Ann E. Walts, Arkadiusz Gertych, Beth Y. Karlan, Alexander M. Xu, Sandra Orsulic

**Author notes:** Corresponding authors: Alexander M. Xu and Sandra Orsulic, Address: David Geffen School of Medicine at UCLA, 10833 Le Conte Ave., 22-177 CHS, Los Angeles, CA 90095. Department of Obstetrics and Gynecology, Tel Aviv Sourasky Medical Center, Tel Aviv 6423906, Israel (Yael Raz). Equal contribution (Maria Sol Recouvreux, Kathleen Trang). **Disclosure:** The authors declare no potential conflicts of interest.

## Abstract

Despite its name, high-grade serous ovarian carcinoma (HGSC) originates in the fallopian tube, not the ovary, arising from a morphologically recognizable precursor lesion, serous tubal intraepithelial carcinoma (STIC). Yet the early cellular and microenvironmental changes driving this transformation remain poorly understood, limiting progress in early detection, interception, and prevention. Here, we generated a Visium HD spatial transcriptomic atlas of fallopian tube carcinogenesis spanning histologically unremarkable fallopian tube epithelium (FTE), STIC, and invasive HGSC. This approach enabled unbiased, tissue-wide, whole-transcriptome mapping at single-cell-level resolution within preserved histologic architecture, providing spatial granularity beyond prior region-of-interest-based platforms. STIC lesions displayed a coordinated epithelial transformation program marked by proliferation, replication stress, DNA repair activation, chromatin remodeling, and induction of tumor-associated antigens, including PRAME and CLDN6, which are emerging targets for vaccine and antigen-directed therapeutic strategies. In contrast, histologically unremarkable FTE contained spatially restricted epithelial defense programs marked by SCGB1A1 and MUC6, suggesting localized protective states that may influence susceptibility to malignant transformation. Using distance-and density-aware spatial analyses, we found that precursor lesions were embedded within immune-enriched, stromal-depleted microenvironments characterized by interferon-dominant immune activation, attenuation of TNFα/NF-κB signaling, macrophage and lymphoid remodeling, and extracellular matrix-associated fibroblast interactions. Computational pathology analysis of collagen architecture confirmed reduced collagen fiber density in STIC-adjacent stroma, linking transcriptomic evidence of stromal remodeling to structural extracellular matrix changes. Together, these data define early epithelial, immune, and stromal programs associated with STIC, and identify tumor-associated antigens, epithelial defense states, and immune-stromal niches as candidate targets for HGSC prevention and early interception.

**CLINICAL RELEVANCE:** A cell-level resolution spatial atlas reveals that ovarian cancer precursor lesions activate vaccine-targetable antigens and remodel their immune-stromal niche well before invasive disease emerges, while normal fallopian tube epithelium relies on previously uncharacterized local defense programs to resist transformation.

## INTRODUCTION

Ovarian high-grade serous carcinoma (HGSC) is one of the most aggressive gynecologic malignancies, accounting for over 20,000 new diagnoses and more than 12,000 deaths annually in the United States ^1^. Most patients with HGSC present with advanced-stage disease, and although tumors often respond initially to cytoreductive surgery and platinum-based chemotherapy, relapse is common. Survival remains poor, particularly for patients with distant-stage disease, for whom 5-year relative survival is approximately 31.5% ^1^. During the past decade, therapeutic advances, including PARP inhibitors, anti-angiogenic therapy, and molecularly guided maintenance strategies, have improved outcomes for selected patients ^2^. However, comparable progress has not been achieved in early detection or primary prevention, particularly for strategies directed at the earliest precursor lesions of HGSC.

Over the past two decades, the dominant model of HGSC pathogenesis has shifted from an ovarian surface epithelial origin toward a fallopian tube origin for many cases ^3–5^. This paradigm emerged largely from detailed pathologic examination of risk-reducing salpingo-oophorectomy specimens from BRCA1/2 pathogenic variant carriers, particularly through the Sectioning and Extensively Examining the Fimbriated End protocol (SEE-FIM), which increases detection of occult lesions in the distal fallopian tube and fimbria ^6, 7^. Together with p53 and Ki-67 immunohistochemistry, careful morphologic assessment has supported a putative histologic continuum of fallopian tube carcinogenesis that includes p53 signatures, serous tubal intraepithelial lesions (STIL), and serous tubal intraepithelial carcinoma (STIC)^8, 9^. p53 signatures are characterized by aberrant p53 staining in morphologically benign fallopian tube epithelium, whereas serous tubal intraepithelial lesions show atypia that falls short of STIC ^8, 9^. STIC lesions exhibit marked cytologic atypia, aberrant p53 expression, and increased proliferative activity, and are widely regarded as immediate precursors to many HGSCs ^4, 8, 10^. This relationship is supported by frequent fimbrial localization and shared TP53 mutations between STIC and concurrent invasive carcinoma, although not all HGSCs have an identifiable STIC and the biological trajectories of STIC progression remain incompletely understood ^4, 10–13^.

The distinction between an incidental STIC and a STIC identified in a patient with concurrent HGSC is also less absolute than these clinical categories imply. Women in whom STIC is identified during risk-reducing surgery remain at substantial subsequent risk of pelvic high-grade serous carcinoma or peritoneal carcinomatosis, whereas the corresponding risk after risk-reducing surgery without STIC is extremely low. Thus, the absence of clinically detectable carcinoma at the time of STIC removal does not necessarily establish complete biological isolation from disseminated microscopic disease below the threshold of clinical or pathologic detection. The possibility of unsampled microscopic tubal carcinoma represents an additional limitation, because even rigorous sectioning cannot exclude disease in tissue that was not examined. Moreover, incidental STICs are often identified in women undergoing prophylactic surgery at younger ages, whereas STICs associated with concurrent HGSC are more commonly identified in older, postmenopausal women. Consequently, observed differences in the STIC microenvironment may reflect not only the influence of concurrent cancer but also age-and menopause-associated changes in the fallopian tube, making these effects difficult to disentangle.

Recent advances in spatial transcriptomics and proteomics have provided unprecedented insights into the molecular mechanisms underlying fallopian tube carcinogenesis. Prior studies, largely using the GeoMx Digital Spatial Profiler, have shown that STIC lesions are associated with localized transcriptional, immune, and microenvironmental changes, including inflammatory signaling, chromosomal instability, and stromal remodeling ^14–19^. These studies established the importance of spatial context, but were generally limited by the coarse resolution of region-of-interest-based analysis. Other spatial transcriptomic strategies have emerged that achieve single-cell resolution using RNA-binding probes or spatially tagged sequencing ^20, 21^. For example, the Visium spatial transcriptomic method uses an array of surface-bound capture areas to localize gene expression to 55 μm spots spaced 100 μm apart ^22^, while the updated Visium HD platform is capable of whole-transcriptome mapping to 2 x 2 μm continuous barcoded squares ^23^.

Here, we present a Visium HD spatial transcriptomic atlas of STIC-associated fallopian tube carcinogenesis spanning 21 capture areas from 18 patients. Unlike region-targeted spatial profiling approaches, Visium HD enables unbiased, sequencing-based spatial whole-transcriptome profiling across intact tissue sections at single-cell resolution. We focused almost exclusively on postmenopausal patients to minimize potential confounding from age-and menopause-associated differences in the fallopian tube microenvironment. The platform uses a continuous array of 2 × 2 μm barcoded squares and permits gene expression data to be overlaid onto matched H&E images, enabling high-resolution molecular mapping within preserved tissue architecture ^23^. This design allowed us to distinguish STIC epithelial compartments from adjacent histologically unremarkable fallopian tube epithelium, invasive HGSC, immune-enriched regions, and stromal microenvironments.

Our dataset captures a spatial continuum from histologically unremarkable fallopian tube epithelium (FTE), through STIC, to invasive HGSC. By integrating histopathology with whole-transcriptome spatial profiling, we mapped gene expression programs in their native tissue context, identified epithelial and microenvironmental states associated with early transformation, and defined candidate interactions between precursor lesions and surrounding stromal and immune compartments. Our study validated previously reported molecular and spatial alterations in STIC and identified additional features of early carcinogenesis that were not readily detectable using lower-resolution or region-selected spatial platforms. To support future studies of the molecular and spatial dynamics of HGSC initiation, we have made the spatial transcriptomic and imaging dataset publicly available through the Gene Expression Omnibus under accession number GSE303022.

## RESULTS

### Visium HD shows that spatial transcriptomic identity mirrors cellular morphology

Visium HD spatial transcriptomics was used to analyze 21 capture areas (6.5 mm x 6.5 mm) registered to FFPE H&E-stained fallopian tube specimens collected from 18 patients. Seventeen patients were postmenopausal, and eight carried pathogenic germline variants associated with increased ovarian cancer risk (**Suppl. Table 1**). Designations of FTE, p53 signatures, STIL, and STIC lesions were based on consensus pathology criteria that combined H&E histology with p53 and Ki67 immunostaining ^9^. All 21 samples were combined into a single dataset for quality control (see Methods, **Supp. Fig. 1A**). After quality control, 62.01% of segmented cells across all 21 samples were retained for downstream annotation and analysis. Cell type annotations across all 21 samples were processed following Harmony batch correction and unsupervised graph clustering with the Louvain method, which identified ten major cell types (T cells, B cells, myeloid, neutrophils, smooth muscle, fibroblasts, vascular endothelial, lymphatic endothelial, secretory epithelial, and ciliated epithelial cells (see Methods, **Supp. Fig. 1B, C**). Harmony-corrected unsupervised clusters were further combined with pathologist-reviewed annotations to classify FTE, STIC, and HGSC epithelial cells. Overall Expression (OE) analysis was used to distinguish T cell and macrophage subtypes by aggregating expression of relevant marker genes. The cell proportions and clinical metadata (hormone replacement therapy (HRT), mutation status, menopausal status, and patient age) for each sample are summarized in **Fig. 1A** and **Supp. Table 1**. Lastly, the spatial connectivity of epithelial cells was used to further define continuous regions of normal FTE enriched in secretory cells and STIC lesions at single-cell resolution. In total, we identified 20 distinct cell types and one unknown cluster, which contained cells with inconsistent gene expression profiles or morphological properties inconsistent with epithelial cell identity. The gene expression profiles of these groups for selected marker genes and OE scores are shown in **Fig. 1B**.

**Fig. 1.**
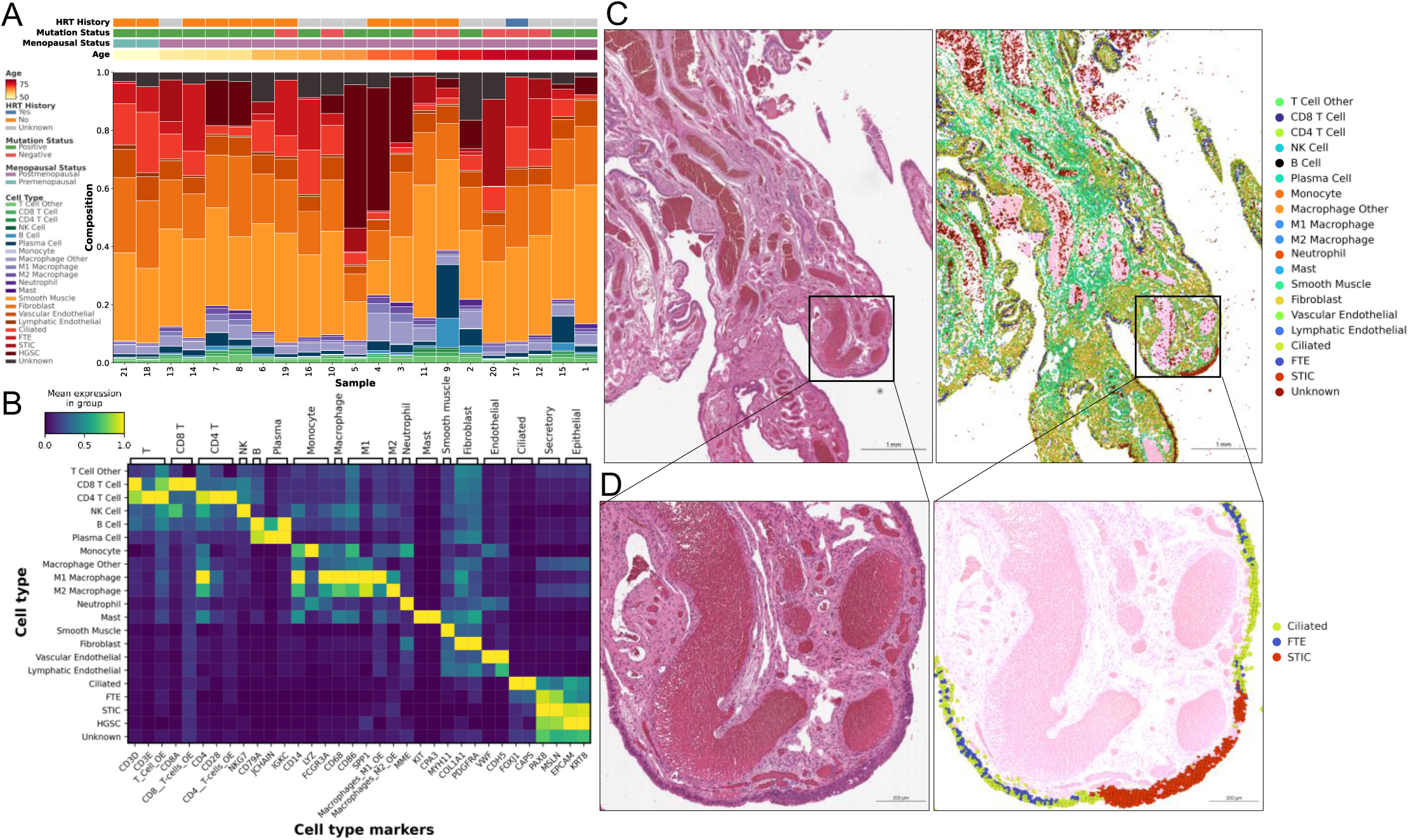
Visium HD spatial transcriptomics of FTE, STIC, and HGSC enables high-resolution, semi-supervised annotation of epithelial, immune, and stromal compartments. **A.** Sample-level cell composition plotted by increasing patient age, with annotated clinical characteristics (full clinical characteristics in Supp. Table 1). **B.** Heatmap showing mean expression of canonical cell-type marker genes across annotated cell groups. Overall expression (OE) scores for xCell gene signatures are shown alongside marker genes for cell types identified using OE-based scoring. **C.** Representative H&E-stained tissue section and corresponding spatial map of annotated cell types. FTE represents histologically unremarkable epithelium computationally enriched for secretory cells by subtracting a curated list of ciliated cell-specific marker genes. **D.** Higher-magnification view of ciliated cells and secretory FTE adjacent to a STIC lesion, with Visium HD overlay demonstrating distinct spatial transcriptional identities.

Cell populations identified by Visium HD were validated through pathologist-guided visual comparison with corresponding cells or cell groups that could be reliably recognized in H&E-stained sections, including epithelial cells, fibroblasts, plasma cells, and immune cell populations phenotypically characterized by small nuclei (**Fig. 1C**). Fallopian tube epithelium contains two major morphologically and transcriptionally distinct epithelial populations, secretory and ciliated cells, which are typically interspersed along the epithelial surface. Examination of overlaid Visium HD transcriptomic expression showed strong spatial alignment between cell annotations and morphological features consistent with secretory or ciliated epithelial identity (**Fig. 1D**).

### STIC lesions display hallmarks of aggressive disease

Spatial analysis of genes differentially expressed between STIC and FTE was limited to 9 STIC lesions in which a visible transition from normal FTE to STIC was present within the capture area (**Fig. 2A**). Two STIC lesions were incidental and 7 were associated with concurrent HGSC. Because normal FTE contains both secretory and ciliated epithelial cells, whereas STIC is thought to arise predominantly from the secretory epithelial lineage and is characterized by marked depletion of ciliated cells ^4, 8, 10^, we defined the FTE cluster primarily by secretory cell identity, with ciliated cells depleted using a curated list of ciliated cell-specific genes derived from our single-cell RNA-seq dataset (GSE298475 and **Suppl. Table 2**). This strategy improved the biological comparability of FTE and STIC by reducing the likelihood that abundant ciliated cell transcripts in FTE would dominate differential expression analysis and obscure transcriptional changes associated with secretory cell transformation.

**Fig. 2.**
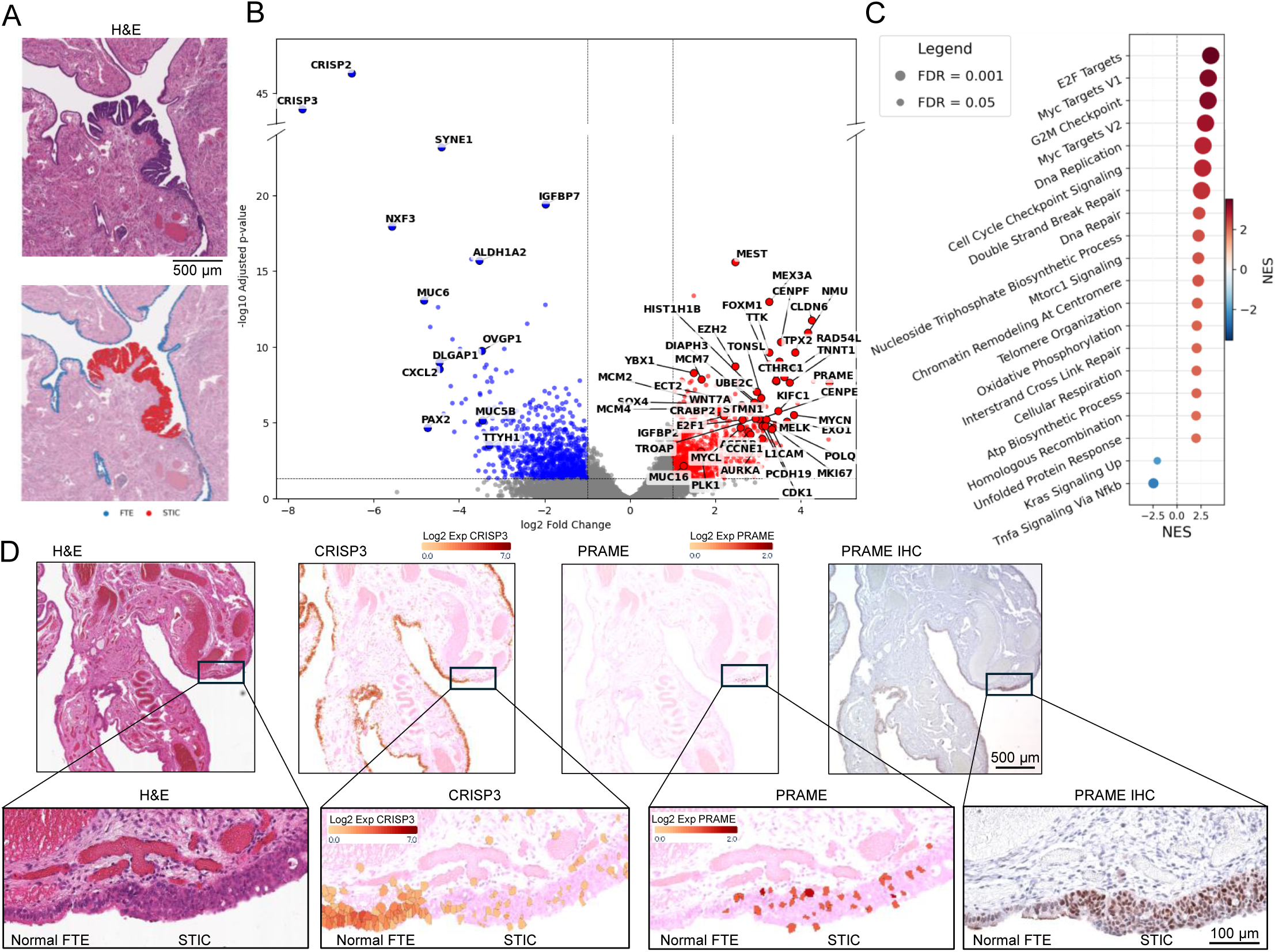
Proliferation is the predominant functional hallmark of STIC lesions. **A.** An example of an H&E-stained section (top) and corresponding Visium HD spatial overlay (bottom) showing cells transcriptomically classified as FTE (blue) or STIC (red). **B.** Volcano plot of differentially expressed genes (DEGs) between FTE (blue) and STIC (red). Ciliated cell-specific genes were excluded from this analysis to reduce bias from the ciliated-cell transcriptomic program, which is more strongly represented in FTE than in STIC. **C.** Gene set enrichment analysis of selected MSigDB Hallmark and Gene Ontology biological process pathways differentially enriched in FTE and STIC lesions. Significant pathways of interest are on the y-axis, while normalized enrichment score (NES) is plotted on the x-axis. Ciliated cell-specific genes were removed from this analysis. **D.** Representative H&E-stained section of the fimbria containing a STIC lesion. The boxed region highlights the transition between FTE and STIC at higher magnification. Shown are Visium HD spatial overlays for CRISP3, a gene downregulated in STIC, and PRAME, a gene upregulated in STIC. IHC also validated PRAME upregulation in STIC.

Genes identified as upregulated in STIC lesions relative to FTE are shown in **Fig. 2B** and **Suppl. Table 3**. Seven upregulated genes (PRAME, UCHL1, TPX2, EZH2, ASF1B, S100A4, and PLK1) were selected for validation based on antibody availability for IHC. These differentially expressed genes, which had adjusted *p*-values ranging from 0.01 to 2.41 × 10⁻¹^0^ and log2 fold changes ranging from 1.67 to 3.88 (**Suppl. Table 3**) were all validated by IHC (**Suppl. Fig. 2**). Genes and pathways upregulated in STIC lesions predominantly reflected increased cell proliferation (**Fig. 2B**, **C** and **Suppl. Table 4**) and were further subdivided into distinct functional categories. First, the dominant signal reflects activation of cell-cycle progression and the mitotic machinery, indicating sustained proliferative activity. Second, this proliferative program is coupled to enhanced DNA replication and replication stress response supported by increased expression of MCM4, GINS1, TICRR, and RRM2 genes involved in elevated origin firing, replication fork progression, and nucleotide biosynthesis. Third, upregulation of BRCA2, RAD54L, EXO1, and POLQ indicates activation of DNA damage response and repair pathways as compensation for replication-associated genomic instability. Fourth, increased expression of FOXM1, MYBL2, E2F1, and EZH2 indicates integration of proliferation, chromatin regulation, and transcriptional amplification, reinforcing the hyperproliferative phenotype. Fifth, elevated levels of UHRF1, ASF1B, HIST1H1B, and HMGA1 suggest dynamic chromatin restructuring that facilitates both DNA replication and transcriptional reprogramming. Sixth, upregulation of KIF11, KIF23, NUSAP1, and PRC1 reflects cytoskeletal reorganization, active microtubule dynamics, spindle stabilization, and cytokinesis, linking structural machinery to mitotic execution. Seventh, increased expression of CLDN6, L1CAM, CTHRC1, and WNT7A suggests early changes in tissue organization and microenvironmental interaction that might support dissemination. Eighth, upregulation of MTHFD2, GLDC, SCD, and IGFBP2 indicates shifts in metabolic reprogramming and growth factor signaling necessary to support biosynthetic and redox demands of rapid growth. Finally, expression of lineage-associated and cancer-testis–related genes, including PRAME, CRABP2, CTCFL, and MEX3A, reflects a dedifferentiated and transcriptionally permissive state.

Notably, this gene set was upregulated in a STIC lesion but not in a STIL lesion from the same patient (**Suppl. Fig. 3**), suggesting that the transition to STIC is not characterized by isolated gene changes, but rather by the emergence of a coordinated proliferative and replication-associated program, marked by upregulation of genes involved in DNA synthesis, mitotic progression, and chromatin regulation, including genes with overlapping or redundant functions. This redundancy may limit the effectiveness of targeting individual genes and instead supports the identification of critical bottlenecks within the proliferative program that are required to sustain high replication rates. Histone chaperones are non-enzymatic regulators of chromatin assembly that can become rate-limiting under oncogene-driven proliferation. ASF1A and its paralog ASF1B function at a key S-phase bottleneck, where newly synthesized DNA must be rapidly assembled into chromatin to maintain replication efficiency while preserving genome integrity. We observed a significant upregulation of ASF1B and a modest increase in the functionally redundant ASF1A in STIC lesions. ASF1B is amplified in more than 10% of HGSC and is associated with poor survival ^24^. To determine whether selective disruption of ASF1B is sufficient to create a vulnerability in ovarian cancer cell proliferation despite potential partial compensation by ASF1A, we targeted ASF1B in the highly aggressive ovarian cancer cell line SKOV3ip1, characterized by intermediate expression levels of both ASF1B and ASF1A. ASF1B-knockout SKOV3ip1 cells exhibited reduced proliferation and invasion in vitro, and formed smaller tumors in vivo compared to control cells (**Suppl. Fig. 4**), supporting a functional role for ASF1B in promoting ovarian cancer proliferation.

### Spatially restricted epithelial defense programs distinguish normal FTE from STIC

Even after selecting FTE areas enriched for secretory cells and removing ciliated cell-specific genes from the analysis (**Suppl. Table 2**), most genes and pathways downregulated in STIC lesions (**Suppl. Table 3**) were predominantly those expressed in ciliated cells, consistent with the reduced abundance of this cell type in STIC lesions ^4, 8, 10^. Notably, several genes normally enriched in secretory FTE were also downregulated, indicating alterations beyond the simple loss of ciliated cells. These included PAX2, one of the first proteins reported to be downregulated in STIC lesions ^25, 26^, CRISP2/3, DLGAP1, TTYH1, IGFBP7, CXCL2, MUC6, SLITRK2, and SYNE1 (**Fig. 2B**). IHC analysis revealed that SYNE1 (nesprin-1) was not only downregulated in STIC lesions but also that this downregulation was accompanied by loss of nuclear envelope localization in pleomorphic STIC epithelial cell nuclei (**Suppl. Fig. 5**).

Pathways significantly enriched in FTE included TNFα signaling through NFκB (**Fig. 2C** and **Suppl. Table 4**). We have previously shown that ciliated cells protect the epithelium by activating TNFα signaling in DNA-damaged secretory cells through the STING paracrine effect which requires functional p53 ^27^. This can activate NFκB, one of the major transcriptional drivers of the senescence-associated secretory phenotype (SASP), consistent with the observed enrichment of IGFBP7 ^28^ and CXCL2 ^29^ in FTE (**Suppl. Table 3**). These findings support a model in which TNFα/NFκB signaling can function as part of an acute senescence-associated barrier, but persistent activation may convert this response into a pro-tumorigenic inflammatory state ^14^.

One of the major strengths of Visium HD compared to GeoMx or conventional laser-capture microdissection is its ability to continuously survey whole-transcriptome expression across the entire tissue section without preselection. This tissue-wide view uncovers focal gene-expression patterns and reveals sublineages of morphologically indistinguishable cells that nonetheless carry distinct transcriptional programs, including higher-order spatial responses to local environmental stress. We found that a subset of genes downregulated in STIC lesions marks spatially confined domains of FTE that would likely be obscured or averaged out using other technologies, underscoring the value of unbiased tissue-wide profiling.

One such focally expressed transcript was SCGB1A1 (uteroglobin, also known as Clara cell secretory protein CC10 or CC16) (**Fig. 3A**). In the airway epithelium, SCGB1A1-positive club cells act as sentinel progenitors that self-renew after injury and generate differentiated ciliated cells to rapidly restore denuded epithelium, a regenerative capacity that has led to their proposed role as progenitor cells for lung cancer ^30–34^. Club-like SCGB1A1-expressing cells have also recently been proposed as prostate cancer progenitor cells based on their identification in proliferative inflammatory atrophy, a lesion associated with prostate cancer precursor states ^35, 36^. Comparable club-like cells have not been identified in the fallopian tube, although increased uteroglobin expression has been reported in the context of tubal infection and ectopic pregnancy ^37^. In our data, SCGB1A1 expression was frequently confined to discrete epithelial patches of both secretory and ciliated cells and often associated with locally disorganized cells (**Fig. 3B**) or focal areas in which individual cells exhibit features of degeneration (**Fig. 3C**). IHC analysis of a separate cohort of 20 non-malignant premenopausal and postmenopausal fallopian tubes confirmed uteroglobin localization to discrete epithelial patches in postmenopausal fallopian tubes with rare expression in premenopausal fallopian tubes (**Fig. 3D, E**).

**Fig. 3.**
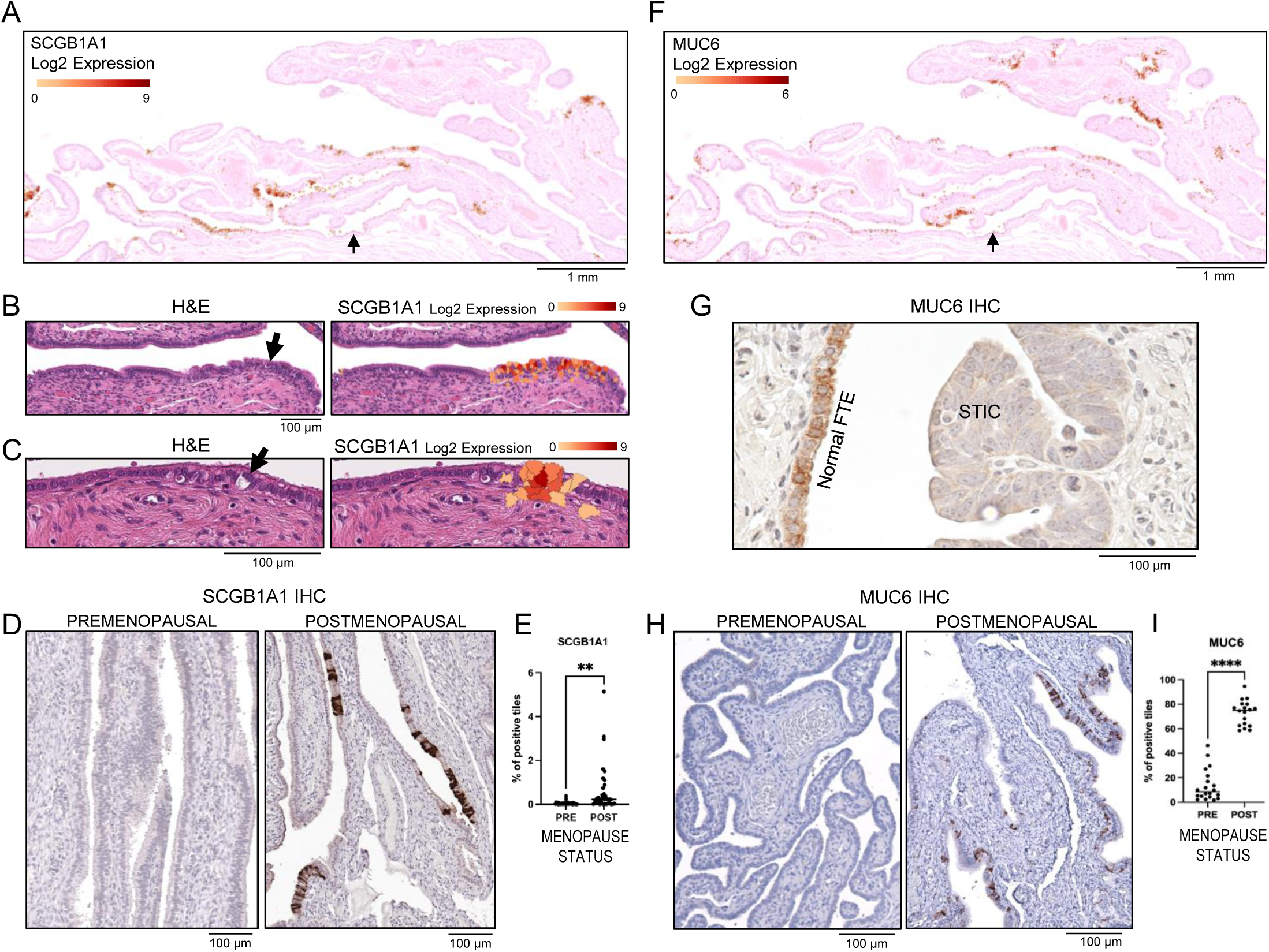
Spatially restricted expression of the mucosal protective genes SCGB1A1 and MUC6 supports localized stress adaptation in FTE. A. Visium HD spatial overlay of SCGB1A1 expression in a fallopian tube fimbria containing an incidental STIL lesion (arrow). B. Regional SCGB1A1 expression in epithelial cells with disrupted structural organization in the fallopian tube (arrow). C. Focal SCGB1A1 expression in a ciliated cell showing degenerative changes (arrow) and in adjacent surrounding cells. D. Representative IHC images showing SCGB1A1 protein expression in a premenopausal and postmenopausal non-malignant fallopian tube. E. Quantification of SCGB1A1 protein expression in a cohort of 10 premenopausal and 10 postmenopausal non-malignant fallopian tubes. QuPath software was used to measure the percentage of SCGB1A1-positive tiles. Statistical significance was assessed using an unpaired t-test; p < 0.01. F. Visium HD spatial overlay of MUC6 expression in a fallopian tube fimbria containing an incidental STIL lesion (arrow). G. Representative IHC image of MUC6 protein expression in a fallopian tube fimbria containing a STIC lesion. H. Representative IHC images showing MUC6 protein expression in a premenopausal and postmenopausal non-malignant fallopian tube. I. Quantification of MUC6 protein expression in a cohort of 10 premenopausal and 10 postmenopausal non-malignant fallopian tubes. QuPath software was used to measure the percentage of MUC6-positive tiles. Statistical significance was assessed using an unpaired t-test; p < 0.0001.

Another transcript identified in discrete regions of FTE was MUC6 (**Fig. 3F**), a gel-forming mucin characteristic of pyloric-type, glandular epithelium that responds to injury by producing a dense protective barrier ^38^. MUC6 was downregulated in STIC lesions, whereas another mucin, MUC16 (CA125) was upregulated in STIC lesions (**Suppl. Table 3**). IHC analysis confirmed MUC6 absence in STIC lesions (**Fig. 3G**) and patchy expression in the FTE (**Fig. 3H, I**).

The focal distribution of SCGB1A1 and MUC6 in FTE is consistent with localized mucosal protection and repair or homeostatic regulation, analogous to their roles in the airway epithelium and pyloric glandular epithelium, respectively. The absence of MUC6 in STIC lesions indicates that this spatially restricted protective program was not maintained during early neoplastic transformation. We hypothesized that expression of SCGB1A1 and MUC6 proteins might be progressively attenuated with aging or reduced hormonal signaling. However, these proteins were unexpectedly expressed primarily in postmenopausal fallopian tubes at both protein (**Fig. 3E, I**) and transcript levels (**Suppl. Fig. 6A-C**). This effect was not due to a global increase in mucin expression, as MUC16 was downregulated in postmenopausal tubes (**Suppl. Fig. 6D**). The reciprocal upregulation of MUC6 and downregulation of MUC16 was validated in an independent bulk RNA-seq dataset of benign fimbrial samples from 24 premenopausal and 48 postmenopausal women (GSE297643) (**Suppl. Fig 6E**).

### STIC and HGSC microenvironments show distance-dependent enrichment of macrophages and T cells and depletion of stromal cells

The distinct transcriptomic signatures of STIC lesions and FTE included immune regulatory-and adhesion-related genes (**Suppl. Table 3**), suggesting that STIC and FTE might differ in their capacity to recruit, maintain, and shape local microenvironments. To define these differences, we analyzed STIC-and FTE-associated microenvironments using four spatial approaches (**Fig. 4A**). First, similar to GeoMx-style analyses, we analyzed cell composition within a 100 µm search radius from FTE or STIC cells (**Fig. 4B-D**). Next, by calculating the density of STIC or FTE cells, we defined new thresholds to consider the influence of density-dependent interactions such as paracrine signaling (**Fig. 4E, F**; see Methods). Third, we quantified radial cell-type distribution to identify fine-grained enrichment and depletion patterns within 10-100 µm in FTE, STIC, and HGSC microenvironments (**Fig. 4G-I**). Finally, we performed epithelial-proximal niche analysis to identify FTE-and STIC-associated microenvironments that influence response to neoplastic epithelial changes (**Fig. 4J, K**).

**Fig. 4.**
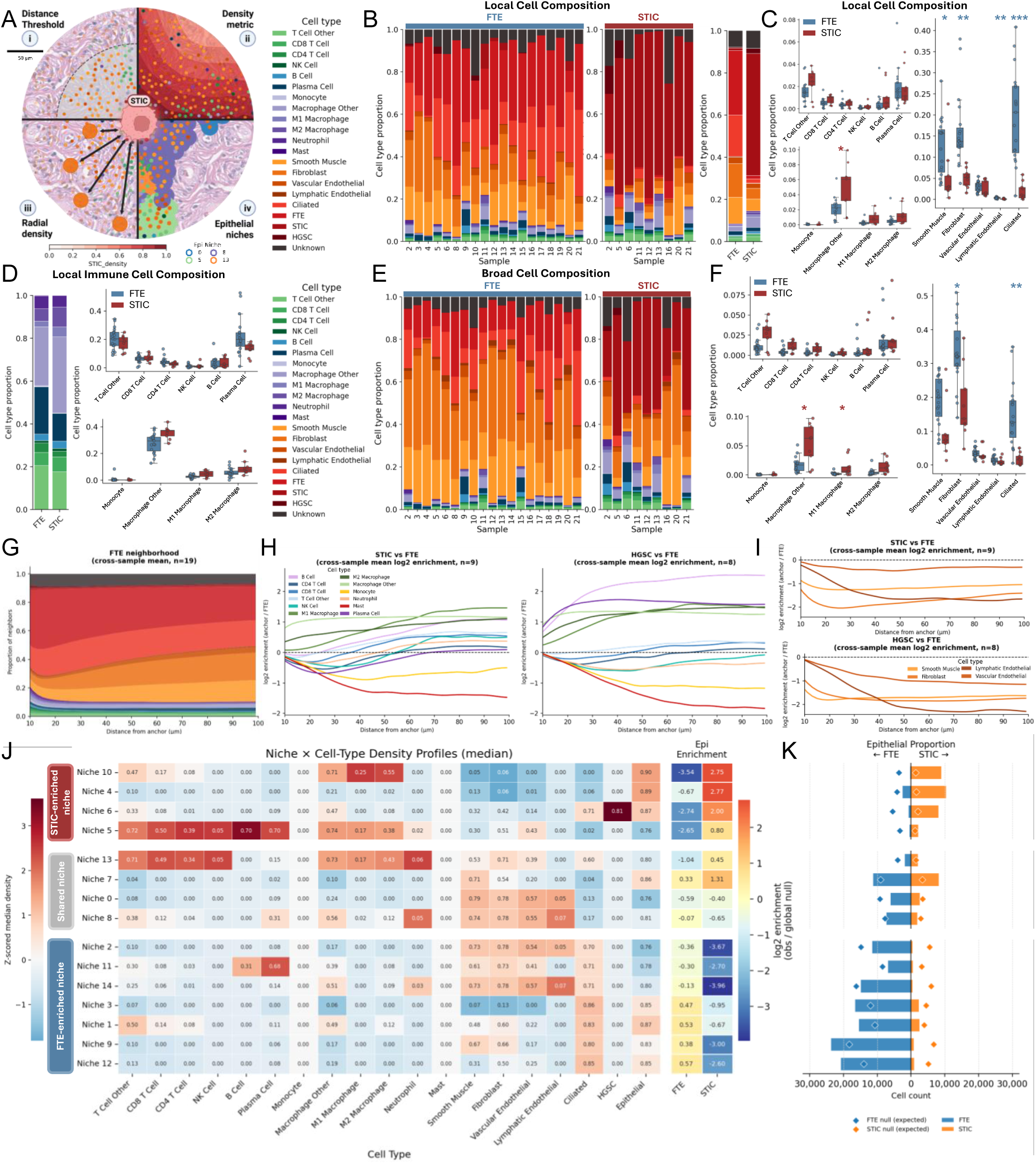
Spatial composition of fallopian tube microenvironments. **A.** Schematic of four spatial metrics used to analyze epithelial-associated microenvironments. These include i. a simple distance-threshold metric identifying cells within 100 µm of the epithelium, analogous to GeoMx-style region-based analyses, used in panels B–D; ii. a nearest-neighbor metric sensitive to epithelial density within 200 µm, used in panel E; iii. radial single-cell distribution analysis across 10–100 µm, used in panels G–I; and iv. epithelial-proximal niche analysis, used in panels J and K. **B.** Cell composition of local FTE- and STIC-associated microenvironments within 100 µm of epithelial cells, shown for each sample and for the combined dataset. **C.** Cell type-specific comparisons of cell proportions in FTE-and STIC-associated microenvironments, stratified by lymphoid, myeloid, and stromal compartments. Mann-Whitney U test; *p < 0.05, **p < 0.01. **D.** Immune cell composition of FTE- and STIC-associated microenvironments within 100 µm, shown across combined samples (left) and immune cell type-specific comparisons expressed as a fraction of total immune cells (right). **E.** CIBERSORTx-assisted analysis of cell composition in broader FTE-and STIC-associated microenvironments within 200 µm using the density threshold. **F.** Cell type-specific comparisons of cell proportions, stratified by lymphoid, myeloid, and stromal compartments. Mann-Whitney U test; *p < 0.05, **p < 0.01. **G.** Radial proportion plot showing the distribution of all cell types surrounding FTE cells across 10–100 µm. **H.** Radial enrichment plots of immune cells surrounding STIC cells and HGSC cells across 10–100 µm. **I.** Radial enrichment plots of stromal cells surrounding STIC cells and HGSC cells across 10–100 µm. **J.** Epithelial-proximal niches defined by cell type density, with niche enrichment for STIC versus FTE. Four niches were STIC-enriched and seven niches were FTE-enriched relative to global expected levels. **K.** Butterfly plot showing the observed counts of FTE and STIC cells and the expected counts of FTE and STIC cells in each niche. Diamonds indicate expected values.

Within a 100 µm radius, STIC and FTE microenvironments differed across three major cellular compartments: immune cells, stromal cells, and ciliated cells. T cells, including CD8^+^ T cells, CD4^+^ T cells, and other T cell subsets, as well as B cells and macrophages, including M1-like, M2-like, and other macrophage populations, were enriched in STIC-associated microenvironments (**Fig. 4B, C**). In contrast, FTE-associated microenvironments contained significantly higher proportions of stromal populations, including fibroblasts, smooth muscle cells, and lymphatic endothelial cells (mean differences: 11%, 6.9%, 0.26%, respectively). Ciliated cells were also significantly more abundant in FTE-associated microenvironments than in STIC-associated microenvironments (mean difference: 17.61%), consistent with the known depletion of ciliated cells during early serous neoplastic transformation ^4, 8, 10^. Lastly, other macrophages were significantly more common in STIC-associated microenvironments (mean difference: 2.15%). When the analysis was restricted to immune cell subsets, STIC-associated microenvironments were dominated by macrophages and contained lower relative proportions of T cells, B cells, and plasma cells compared with FTE-associated immune microenvironments (**Fig. 4D**).

The spatial resolution of Visium HD enables flexible interrogation of epithelial and stromal neighborhoods. Although the local STIC microenvironment within 100 µm was enriched for immune cells, the adjacent stroma is also likely to contribute to early neoplastic progression. However, stromal regions beyond 100 µm from the epithelium consistently generated lower-quality signals, with 79.61% of cells that failed quality control located more than 100 µm from the nearest epithelial cell. This technical limitation reduced confidence in direct single-cell-level assessment of distal stromal contributions. To recover information from lower-confidence stromal regions that could interact with STIC or FTE cells, we identified spatial regions classified as STIC-or FTE-associated using the density threshold, pseudobulked transcriptomic measurements from cells within these regions that failed quality control, inferred stromal cell proportions within this population using CIBERSORTx ^39^, and integrated these inferred populations with high-confidence cells that passed quality control in STIC-associated microenvironments (**Fig. 4A, E**). Using this expanded strategy at a 200 µm interaction radius to calculate density, the broader STIC-associated microenvironment showed higher median proportions of immune cells, including T cells, B cells, plasma cells, and macrophages. Conversely, the broader FTE-associated microenvironment was significantly enriched for ciliated cells and fibroblasts compared with STIC-associated regions, mirroring the differences observed in the local epithelial microenvironment (**Fig. 4B**). Together, these analyses suggest that spatial variation in data quality can be partially addressed by combining single-cell-resolution analysis in high-confidence regions with pseudobulk inference across lower-confidence stromal areas.

After first defining the overall cellular composition of STIC-and FTE-associated microenvironments independent of distance-dependent effects, we next quantified cell distributions as a function of radial distance from the epithelial compartment. Using 10–100 µm as a continuous spatial axis, we compared STIC-and HGSC-associated microenvironments with FTE as the baseline reference (**Fig. 4A, G**). The most significant differences in immune cell radial distributions included enrichment of macrophages and B cells, together with reduced representation of mast cells and monocytes, in both STIC-and HGSC-associated microenvironments relative to FTE (**Fig. 4H**). Several immune populations showed distance-dependent patterns that differed between STIC and HGSC. HGSC-associated microenvironments were enriched for plasma cells and depleted for NK cells and neutrophils across radial distance. In contrast, radial enrichment of other T cells was unique to STIC-associated microenvironments. CD4^+^ T cells and plasma cells were locally reduced in STIC-associated microenvironments compared with FTE, a pattern not observed in HGSC. Additional immune populations showed radial enrichment patterns specific to STIC-associated microenvironments (**Fig. 4H**).

A pattern not captured by threshold-or density-based analyses was observed for CD8^+^ T cells, neutrophils, and NK cells. These populations showed a radial inversion pattern, characterized by relative depletion near STIC lesions at shorter distances, followed by increased abundance at greater distances. All stromal cell populations, including smooth muscle cells, fibroblasts, lymphatic endothelial cells, and vascular endothelial cells, were radially depleted in both STIC-and HGSC-associated microenvironments compared with FTE, with more pronounced depletion in HGSC-associated regions (**Fig. 4I**). Together, these findings indicate that STIC is associated with early, spatially organized immune remodeling and stromal loss, whereas HGSC exhibits a more advanced immune-restructured and stromal-depleted microenvironment.

The transition from FTE to STIC may precede, accompany, or initiate changes in the local microenvironment. While reciprocal feedback between FTE, STIC, and the surrounding microenvironment is expected, we sought to decouple epithelial state from microenvironmental context by identifying epithelial-proximal niches that were either unique to STIC, unique to FTE, or shared by both. We hypothesized that STIC-specific microenvironments were more likely to reflect microenvironments remodeled by STIC lesions, whereas niches shared by STIC and FTE might represent permissive states that are not specific to early neoplastic transformation. To define these microenvironments, we first grouped FTE and STIC cells into a single epithelial category and filtered the dataset to include cells with an epithelial density ≥ 0.5, thereby focusing on epithelial-proximal microenvironments (see Methods, **Fig. 4A**). Next, we generated spatial niches using only these epithelial-proximal cells and classified niches as STIC-or FTE-enriched niches based on absolute enrichment score difference > 2 (**Fig. 4J, K)**. STIC-enriched niches (Niches 4, 5, 6, 10) were characterized by lower stromal densities and higher immune cell densities in either macrophage-enriched contexts, as seen in Niche 10, or macrophage-and lymphoid-enriched contexts, as seen in Niche 5. In contrast, FTE-enriched niches displayed higher ciliated cell and stromal densities, lower immune cell densities, and lower immune co-localization relative to STIC-enriched niches. B and plasma cells were concentrated in two niches (Niches 5 and 11) that differed in their spatial context. In STIC-enriched Niche 5, B and plasma cells were associated with other immune populations, whereas in FTE-enriched Niche 11, they were associated with stromal-rich environments. Four niches were not enriched in either STIC or FTE, including relatively neutrophil-enriched niches and stromal niches lacking ciliated cells. Together, these findings suggest that STIC lesions are associated with a specific profile of localized immune-rich, stromal-depleted microenvironments, while FTE is associated with ciliated and stromal-rich epithelial neighborhoods.

### Spatially resolved ligand-receptor signaling identifies STIC-associated immune and stromal signaling programs

The largest cellular composition differences between STIC-and FTE-associated microenvironments involved stromal and immune cell proportions. Because fibroblasts, macrophages, and T cells (hereafter defined as target cells) have been implicated in epithelial homeostasis, immune surveillance, stromal remodeling, and early serous neoplastic progression, we next examined spatially-restricted signaling interactions between these target cells and epithelial subtypes using a spatially-subsetted ligand-receptor permutation test. We first identified target cells located near epithelial subtypes, defined by an FTE or STIC density metric ≥0.5 (see Methods). We also identified FTE and STIC epithelial cells located near each target cell, defined by STIC/FTE cells with a target cell density metric ≥0.5 plus STIC/FTE density ≥0.5. We then performed two complementary permutation tests to assess directionally distinct interaction patterns. First, by fixing epithelial cells and permuting the target cell with all other microenvironment cells, we evaluated the significance of the target cell as the signaling partner to the epithelium. Second, by fixing the target cell and permuting all epithelial cells, we evaluated potential changes in the local epithelium induced by the target cell (**Fig. 5A**). These two permutation tests are shown as ligand-receptor interaction arrows originating either from the epithelium or from cell types of interest, respectively (**Fig. 5B**).

**Fig. 5.**
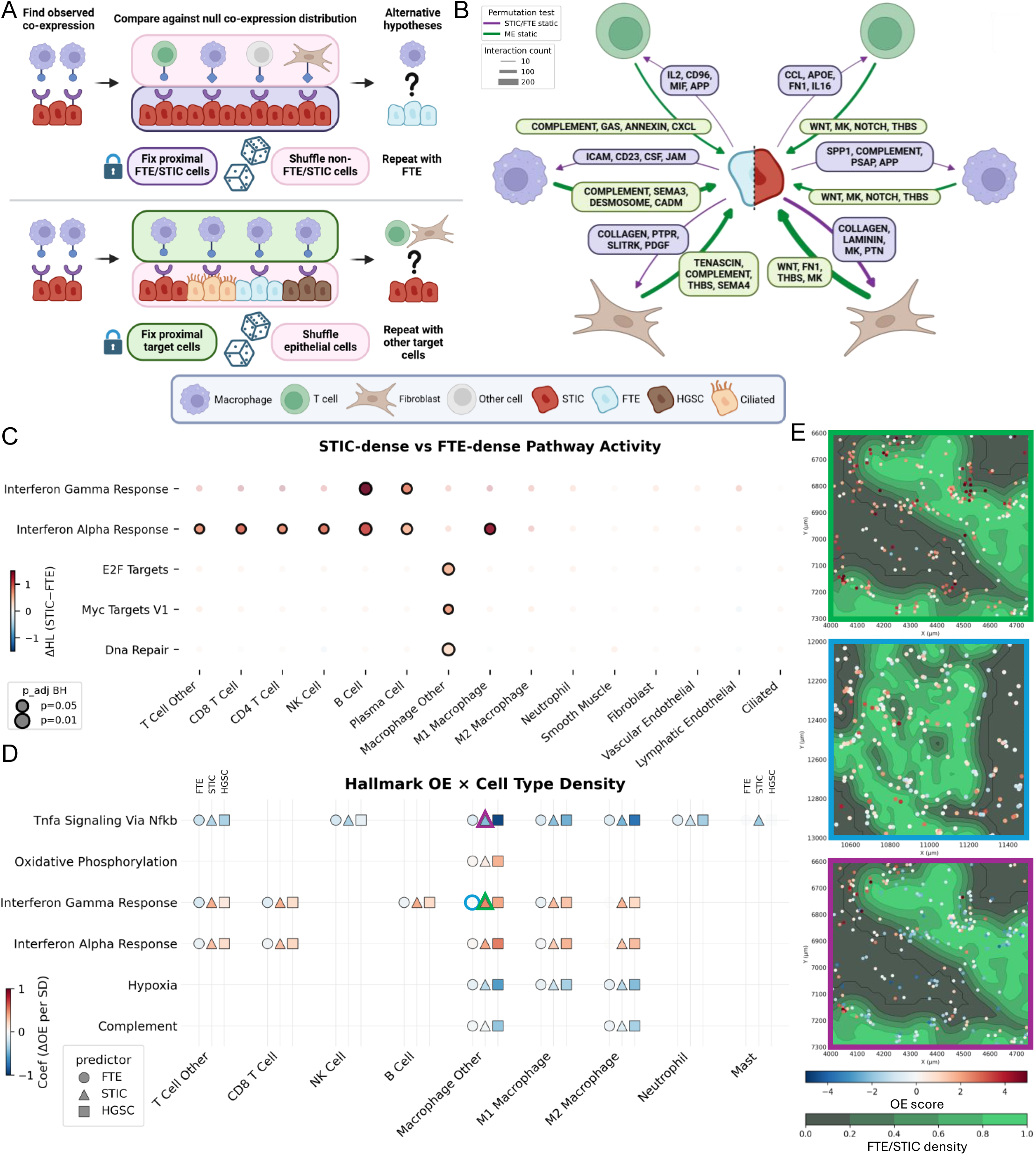
STIC-associated microenvironments exhibit spatially coordinated immune signaling, fibroblast remodeling, and inflammatory pathway activity. **A.** Schematic of epithelial-static and target-cell-static permutation tests used to evaluate spatially restricted ligand-receptor interactions. Observed ligand-receptor co-expression was first calculated between cell types of interest. Null co-expression distributions were then generated by shuffling selected cell labels, and tests were repeated under alternative hypotheses to determine directionally distinct signaling relationships. **B.** Network plot summarizing results from the permutation-based ligand-receptor analysis. Edge weights indicate the difference in significant interactions between STIC-and FTE-associated cells. Each edge is labeled with the top four interacting CellChatDB pathways. ME – microenvironment. **C.** Median differences between cell-specific overall expression (OE) pathway scores between STIC-dense and FTE-dense regions. Color indicates Hodges-Lehmann shifts that estimate median differences in pathway activity between STIC-dense and FTE-dense microenvironments. Only significant differences are shown, p < 0.05. **D.** Coefficients from mixed linear model fitting OEs pathway scores as a function of FTE, STIC, and HGSC density for each cell type. Color indicates the difference in OE score per standard deviation increase in FTE, STIC, and HGSC density. Displayed pathways had at least one relationship with FTE, STIC, or HGSC density with a coefficient > 0.3. **E.** Representative examples showing relationships of STIC/FTE density (contour) with other macrophage OE scores showing a positive correlation (green border, STIC: IFNγ), no correlation (blue, FTE: IFNγ), and negative correlation (purple, STIC: TNFα).

The first analysis with fixed epithelial cells showed that unique STIC-T cell interactions were characterized by a dominant CCL5 chemokine-feedback axis and IL16/CD4 recruitment cue that suggests spatially-organized immune surveillance in STIC microenvironments. Conversely, FTE-T cell interactions were largely defined by IL2 (IL7/IL7R and TSLP/IL7R) and CD96 signaling pathways (CD96/PVR and CD96/NECTIN1) in conjunction with MADCAM1/ITGA4 signaling, consistent with homeostatic tissue surveillance. STIC-macrophage interactions displayed a tumor-associated macrophage (TAM)-like signature, with predominantly SPP1 and complement pathway signaling, suggesting an immunosuppressive myeloid context. Similar to FTE-T cell signaling, FTE-macrophage signaling implies homeostatic immune surveillance with dominant ICAM and CSF signaling pathways (CSF1-CSF1R and IL34/CSF1R). These interactions suggest that immune cells in FTE-associated microenvironments may participate in epithelial maintenance, matrix organization, and immune surveillance. In contrast, fibroblasts interacted more frequently with STIC than with FTE cells through collagen, laminin, and midkine (MK) signaling pathways. MK interactions with ITGA and ITGB subunits are consistent with stromal remodeling in the STIC-associated niche and possible emergence of a cancer-associated fibroblast-like phenotype.

In the complementary fixed-target-cell analysis, STIC cells showed stronger inferred signaling to their surrounding microenvironment than FTE cells, primarily through laminin and collagen pathways. STIC-exclusive T cell and macrophage interactions were dominated by WNT7A-or WNT6-mediated activation of STIC Frizzled receptors (FZD) and MK signaling, suggesting a tumor-permissive immune microenvironment. STIC-exclusive fibroblast interactions occurred through the WNT (WNT7A, WNT10A, WNT6), THBS, and MK signaling pathways. In contrast, FTE-exclusive interactions between immune cells and fibroblasts align with previously implied immune surveillance and epithelial architecture preservation programs through the GAS, complement, tenascin, and THBS signaling pathways. Together, these findings suggest that STIC-associated epithelial cells and fibroblasts engage in reciprocal ECM-related signaling. The enrichment of collagen, laminin, WNT, and MK pathway interactions is consistent with early stromal remodeling, although additional validation will be required to determine whether these signals reflect the emergence of a cancer-associated fibroblast phenotype.

Due to the transcriptomic sparsity of our dataset, to identify coordinated pathway regulation within FTE-, STIC-, or HGSC-dense microenvironments, we used Overall Expression (OE) scores (see Methods) to quantify pathway activity for MSigDB Hallmark gene sets (**Fig. 5C**). T cells, B cells, plasma cells, and M1 macrophages in STIC-dense microenvironments exhibited significantly higher interferon response activity compared with the same cell types in FTE-dense microenvironments. Interferon Alpha (IFNα) Response OE activity was significantly higher in STIC-dense immune cells across major lymphoid and myeloid populations, including other T cells, CD8^+^ T cells, CD4^+^ T cells, NK cells, B cells, plasma cells, and M1 macrophages. This increase in IFNα OE activity was coupled with a significant increase in Interferon Gamma (IFNγ) Response in STIC-dense B cells and plasma cells. In addition, macrophages in STIC-dense microenvironments exhibited significantly higher OE activity for E2F targets, Myc targets V1, and DNA Repair pathways compared to macrophages in FTE-dense microenvironments, suggesting that STIC-associated macrophages may occupy a more activated or proliferative transcriptional state.

After identifying pathway activity differences between STIC-and FTE-dense microenvironments, we sought to determine whether these activity differences were dependent on spatial proximity to FTE, STIC, or HGSC. To address this, we modeled OE scores as a function of FTE, STIC, and HGSC cell density across all immune cell types, using a mixed linear model. This approach enabled detection of density-dependent spatial relationships that could not be resolved by region-level spatial analyses (**Fig. 5D**). For example, macrophages located at progressively greater distances from epithelial regions (**Fig. 5E**, green box), expressed correspondingly greater IFNγ pathway activity only in STIC- or HGSC-associated microenvironments, while distance from FTE showed no comparable relationship (blue box). We found that an increase in IFNγ and IFNα response activity in macrophages, plasma cells, B cells, and T cells was significantly associated with increased STIC density, while IFNγ response was inversely related to increasing FTE density. In plasma cells, B cells, and T cells, the relationship of INFγ and INFα response activity and HGSC density was weaker than the association observed with STIC density, whereas macrophage IFNγ and IFNα response activity increased with both STIC and HGSC density. We observed an opposing pattern for the TNFα/NF-κB pathway. TNFα/NF-κB OE activity was significantly decreased in other T cells, NK cells, and all myeloid cell populations as epithelial density increased across microenvironments (purple box). The magnitude of this decrease followed a disease progression gradient, with the strongest attenuation in HGSC-associated regions, followed by STIC-associated and then FTE-associated regions. Hypoxia pathway activity in macrophages and complement pathway activity in other macrophages and M2 macrophages exemplified a similar pattern to TNFα, while oxidative phosphorylation activity increased along the disease progression-associated density axis. Together, these analyses integrate gene-level expression into pathway-level OE scores and quantify their spatial dependence on epithelial compartment density. The results reveal spatially structured immune signaling programs across early tumorigenesis and progression, with STIC-associated microenvironments showing consistent similarities to HGSC-associated microenvironments and clear differences from FTE-associated microenvironments.

### STIC-adjacent stroma shows reduced collagen fiber density

Differential gene expression analysis of STIC- versus FTE-associated stromal regions revealed enrichment of three extracellular matrix–related genes in STIC-associated stroma: matrix Gla protein (MGP), versican (VCAN), and biglycan (BGN) (**Suppl. Fig. 7A**), which was recently reported to be upregulated in STIC-associated stroma ^40^. Because MGP, VCAN, and BGN are involved in collagen remodeling, fibril organization, and extracellular matrix regulation ^41–43^, and because we previously showed that stromal architecture changes progressively with increasing proximity to STIC lesions ^44^, we next focused on collagen architecture. Masson’s trichrome staining showed subtle differences in collagen distribution and staining intensity between STIC-adjacent and FTE-adjacent stroma, along with increased immune cell infiltrate near STIC lesions (**Suppl. Fig. 7B**). However, these differences were not readily amenable to reproducible visual scoring. To quantitatively assess collagen architecture, we performed computational pathology analysis of annotated STIC-adjacent and FTE-adjacent stromal regions in H&E-stained sections from the same 9 capture areas used to identify differentially expressed genes between FTE and STIC. We quantified median fiber length, the standard deviation of fiber length, fiber density defined as the ratio of fiber pixels to total pixels within each region of interest (ROI) (**Suppl. Fig. 7C**), and fiber orientation entropy ^45^. A total of 58 STIC-adjacent and 61 FTE-adjacent non-overlapping stromal ROIs were analyzed. For each capture area, feature values were summarized as the median across ROIs of the same tissue type within that slide. Of the four features examined, only fiber density (FibDens) was statistically significant (**Suppl. Fig. 7D**). Paired per-case comparisons of median FibDens values between STIC-adjacent and FTE-adjacent stromal ROIs showed a consistent reduction in collagen fiber density in STIC-adjacent stroma across all nine cases (**Suppl. Fig. 7E**). The effect size showed moderate between-case heterogeneity (I² = 53%) (**Suppl. Fig. 7F**).

In the validation cohort, we analyzed 365 HGSC-adjacent, 599 STIC-adjacent, and 630 FTE-adjacent non-overlapping stromal ROIs, corresponding to 20 HGSC, 48 STIC, and 34 FTE stromal regions across 54 cases (**Suppl. Fig. 7G**). Linear mixed-effects modeling showed that fiber density was significantly lower in STIC-adjacent stroma than in FTE-adjacent stroma (β₁ = mean[STIC] − mean[FTE] = −0.0143, p = 3.12 × 10⁻⁵), and even lower in HGSC-adjacent stroma than in FTE-adjacent stroma (β₂ = mean[HGSC] − mean[FTE] = −0.0253, p = 3.05 × 10⁻⁷), indicating a progressive reduction in fiber density from FTE to STIC to HGSC. Variance decomposition revealed substantial between-case variability (σ²_case_ = 3.41 × 10⁻⁴) relative to within-case, ROI-level variability (σ²_ROI_ = 1.83 × 10⁻⁴), corresponding to an intraclass correlation coefficient of ICC = 65.1%. Thus, approximately 65% of the total variability in FibDens was attributable to differences between cases, indicating that case identity is a major determinant of this feature and supporting its sensitivity to biologically meaningful heterogeneity. The model showed good fit (AIC = −498.4), supporting the contribution of tissue context, including FTE-, STIC-, and HGSC-adjacent regions, to variation in fiber density. As a complementary exploratory analysis, per-case median FibDens values were also assessed using the Mann–Whitney U test (**Suppl. Fig. 7G**).

## DISCUSSION

Despite recent breakthroughs in STIC lesion profiling, many fundamental questions about STIC biology, epithelial transformation, and microenvironmental remodeling remain unresolved, limiting progress in early detection, interception, and prevention. To address these gaps, we applied high-resolution spatial transcriptomic profiling, which differs from recent studies by preserving the histologic context of rare STIC lesions and directly linking gene-expression programs to specific epithelial and stromal compartments within the same tissue section.

Our spatial differential gene-expression analyses comparing STIC lesions with FTE are consistent with prior molecular, single-cell, spatial, and proteomic studies ^5, 14, 15, 17, 46, 47^, showing that STIC lesions have already acquired a coordinated transcriptional program characteristic of aggressive HGSC, marked by accelerated proliferation, replication stress, DNA repair activation, chromatin remodeling, and early changes in tissue organization. The absence of this program in a paired STIL lesion suggests that progression to STIC reflects a discrete biological transition rather than a gradual accumulation of isolated gene-expression changes. Because STIC lesions already display advanced molecular features of aggressive HGSC and may therefore be difficult to eradicate once established, greater attention should be directed toward prevention and toward defining the protective mechanisms that preserve normal fallopian tube epithelial homeostasis and prevent transformation ^27^.

Serous carcinogenesis is thought to emerge in a setting of recurrent epithelial stress and injury ^48^ suggesting that the fallopian tube epithelium depends on localized protective programs to preserve tissue integrity in the face of mechanical stress, inflammation, oxidative damage, and other microenvironmental insults. Our study revealed regional, mostly non-overlapping, expression of SCGB1A1 and MUC6, two genes potentially involved in epithelial maintenance and homeostasis. Although the exact functions of SCGB1A1 and MUC6 in the fallopian tube have not been defined, prior associations between their expression and inflammatory conditions in the fallopian tube ^37^, together with their established anti-inflammatory and epithelial repair functions in other epithelial tissues, suggest their involvement in fallopian tube damage control.

SCGB1A1 is primarily produced by airway club cells, where it plays a key role in tissue regeneration after injury and infection ^30, 31^, while MUC6 is characteristic of pyloric-type epithelium, where it contributes to the gastric mucus barrier that protects the epithelium from luminal and microbial stress ^38^. In the stomach, loss of SCGB1A1 and MUC6 has been associated with increased susceptibility to inflammation and carcinogenic progression, supporting the idea that SCGB1A1 and MUC6 may mark a protective epithelial phenotype rather than merely a differentiation state ^49–51^. Together, these observations suggest that SCGB1A1 and MUC6 may be induced in stressed fallopian tube epithelium but act through distinct mechanisms, with SCGB1A1 modulating inflammation and stabilizing repair, and MUC6 providing a physical barrier. Their non-overlapping regional expression further suggests that morphologically normal fallopian tube epithelium is biologically heterogeneous, with selected areas existing in distinct protective states. This concept is clinically important because it suggests that the cell-of-origin landscape is heterogeneous even before morphologic transformation becomes apparent. In this framework, SCGB1A1-positive and MUC6-positive regions may mark territories of active epithelial defense, whereas disruption of these localized stress-response programs may identify areas at greater risk for progression to STIC and invasive carcinoma. More broadly, these findings raise the possibility that regional stress-adaptation programs, rather than a uniform field effect, help shape susceptibility to malignant transformation in the fallopian tube.

Our study also suggests potential preventive strategies. Recombinant human SCGB1A1 (rhCC10) is a small protein amenable to therapeutic development because of its resistance to proteolysis and stability across a broad range of temperatures and pH conditions ^52^. In premature infants with respiratory distress syndrome, early administration of a single dose of rhCC10 was associated with lower leukocyte and neutrophil counts in tracheal aspirates than placebo, supporting its anti-inflammatory activity ^53^.

Given the current lack of effective primary prevention strategies, interception at the STIC stage may now represent a more realistic clinical opportunity. This is especially relevant when STIC lesions are found incidentally in prophylactically removed fallopian tubes, as approximately 20% of affected women develop ovarian cancer within 5 to 10 years of follow-up ^54–56^. This pattern suggests that exfoliated STIC cells have, in some cases, already disseminated to the peritoneal surfaces before surgery. Yet clinical management after incidental STIC detection remains poorly defined, with current options ranging from exploratory laparoscopy and liquid biopsy to empiric chemotherapy or close surveillance ^57^. Our identification of CLDN6 and PRAME upregulation in STIC lesions points to a possible molecularly guided interception strategy. CLDN6 and PRAME are tumor-associated antigens with limited expression in normal adult tissues. CLDN6 is overexpressed in a subset of ovarian cancers relative to benign tissue ^58^, while PRAME is an embryonic antigen that is largely silenced in adult tissues except the testis but is reactivated in malignancy ^59^. Notably, both genes appear to be upregulated in STIC but not in p53 signatures or STIL lesions, suggesting that their induction is linked to the transition to a bona fide preinvasive neoplastic state. Given that both proteins have already been explored as vaccine targets in preclinical and early clinical settings ^58, 60–66^, they could be quickly repurposed as candidates for immune interception at early stages of ovarian cancer development. In this context, antigen-specific vaccination against CLDN6 and PRAME could, in principle, eliminate occult precursor or disseminated cells before invasive disease becomes clinically apparent, offering a path toward precise immunologic interception of ovarian cancer at its earliest detectable stage.

For translational strategies focused on the microenvironment rather than STIC-specific signatures, spatial transcriptomics provides our clearest view to date of how STIC lesions interact bidirectionally with their local microenvironment relative to FTE and HGSC. Across complementary analyses of cell composition, radial proportion distributions, ligand-receptor interactions, and pathway activity scoring, STIC microenvironments appeared immune-enriched and stromal-depleted, with a chronic pro-inflammatory immune response dominated by IFN signaling and tumor-promoting extracellular matrix remodeling involving fibroblast-collagen interactions. Recent GeoMx-based spatial profiling of fallopian tube precursor lesions of HGSC has provided an important framework for understanding STIC progression, highlighting localized IFN signaling, chromosomal instability, and a transition from immune surveillance to immune suppression along the disease progression axis ^14, 17, 18^. Our data complement this work while addressing an additional spatial scale, supporting cell-type annotations in a distance-aware context that preserves local tissue architecture. This allowed us to ask not only whether STIC-rich regions were different from FTE-rich regions, but whether immune cell behavior changed continuously with local FTE, STIC, or HGSC density.

This density-aware analysis revealed a central feature of the STIC microenvironment: immune cells near STIC adopt an IFN-dominant state, with increased IFNγ response activity across lymphoid and myeloid populations in STIC-dense regions and increased IFNα activity in B cells, plasma cells, and macrophages. IFN activity was inversely associated with FTE density, suggesting that the STIC microenvironment represents an early tumor-associated inflammatory state. Uniquely, the increase in IFN-driven inflammatory activity was not accompanied by an increase in TNFα/NF-κB pathway activity, which was attenuated in T, NK, and myeloid cell populations, with the strongest decrease associated with HGSC density, followed by STIC, then FTE. The immune response along the disease progression axis appears to be selective for IFN programs whereas TNFα/NF-κB activity is progressively restrained.

Overall, our study refines the model of STIC progression by identifying disease-associated immune responses in a spatially resolved cellular context. By resolving transcriptional changes as a function of local epithelial density, our data indicate that STIC lesions are associated with a structured immune niche characterized by IFN activation, attenuation of TNFα/NF-κB signaling, stromal depletion, and ECM remodeling. In addition to the finer single-cell resolution afforded by the spatial data, Visium HD also retained useful transcriptomic information beyond high-confidence segmented cells, allowing us to augment single-cell-level cell proportion measurements with pseudobulked measurements from cells that failed QC but remained near FTE and STIC lesions. The use of spatially continuous metrics adds further nuance to spatial interpretation. All spatial omics data are currently generated using static images, which reflect a snapshot in time. By tracking distance-dependent variation, we provide a descriptive framework that is quantitative and reproducible, and can inform probabilistic or biostatistical models as we consider not just the cells that were interacting when the tissue was fixed, but also the cells most likely to interact in a living tissue. This prospective view will be essential for predictive modeling of the tissue response to therapeutic candidates.

Our transcriptomic analyses are complemented by computational pathology analyses of extracellular matrix architecture, providing an orthogonal assessment of stromal remodeling. The finding that collagen fiber density is reduced in STIC-adjacent stroma suggests that stromal remodeling begins at the precursor stage of HGSC. This is consistent with prior second harmonic generation (SHG)-based studies showing that collagen architecture can distinguish normal fallopian tube, p53 signature/STIC precursor lesions, and high-grade serous carcinoma, supporting the concept that collagen remodeling begins before overt invasive disease ^67^. This supports a model in which early neoplastic epithelium communicates with the local microenvironment, potentially through inflammatory, proteolytic, or fibroblast-mediated mechanisms. One possible explanation is that increased macrophage infiltration in the stroma adjacent to STIC contributes to local matrix remodeling by secreting matrix-degrading enzymes, inflammatory mediators, and factors that alter fibroblast function. Alternatively, macrophage infiltration may physically intercalate between collagen fibers, increasing the cellular fraction of the stroma and creating the appearance of reduced fiber density. Because collagen density alone cannot define the underlying mechanism, future integration with spatial transcriptomics, ECM proteomics, matrix metalloproteinase activity, macrophage markers, and fibroblast-state markers will be needed to determine whether this stromal phenotype reflects active collagen degradation, impaired collagen synthesis, immune-mediated remodeling, altered stromal cellularity, or a distinct precursor-associated stromal program.

### Limitations

A core limitation of spatial transcriptomic studies is the variable transcriptomic data quality inherent to profiling archival FFPE tissue. This was the major source of data loss in our study, as a subset of sections did not yield sufficient high-quality transcriptomic data after QC. Although recently preserved FFPE tissue is preferable for spatial transcriptomic analysis, this requirement is difficult to meet for STIC lesions, which are uncommon and often available only retrospectively from archival pathology material. As a result, our cohort necessarily included sections spanning a broad range of specimen ages and associated RNA degradation. We observed that recently preserved samples, particularly those less than one year old, generated the highest-quality transcriptomic data. For integrated analyses across the cohort, we therefore applied normalization and QC strategies to mitigate variability related to RNA quality and slide age.

Single-cell spatial transcriptomic methods such as Visium HD also depend heavily on cell segmentation algorithms to define cell boundaries and assign transcripts accurately. Although Visium HD resolves transcripts to 2 × 2 μm bins, we observed some incorrect assignments, including doublets and imprecise or incomplete cell boundary detection. Segmentation remains particularly challenging in densely packed tissue regions and in areas where different cell types overlap within the section plane. These limitations precluded direct single-cell comparisons of cells within the FTE and STIC microenvironments. Most instances of improper segmentation occurred in stromal regions and were excluded through stringent QC and visual inspection. After cell-type annotation, both annotations and transcripts of interest were visually reviewed to confirm biologically plausible and histologically consistent expression patterns and cell-type organization. However, cells that failed QC still retained informative transcriptomic signatures. To preserve this information, we used a multilevel analytical strategy in which transcriptomic profiles from QC-failing cells were pseudobulked, similar to bulk RNA-seq analysis, to estimate cell-type proportions and perform ligand-receptor analysis. Future studies may further refine this approach by applying spot-like deconvolution strategies, analogous to standard Visium analysis, to cells that do not meet single-cell QC criteria.

## METHODS

### Patient cohorts

The Visium HD spatial transcriptomic dataset comprised 21 fallopian tube specimens (**Fig. 1A**) collected from 18 patients (**Suppl. Table 1**, GSE303022, UCLA IRB#24-000966). Seventeen of the 18 patients were postmenopausal. Eight women harbored germline pathogenic variants in genes related to DNA repair and genomic instability, including BRCA1, BRCA2, CHEK2, CDK12, PALB2, BRIP1, and ATM, while 10 patients were confirmed noncarriers or had a mutation of unknown significance (VUS) (**Suppl. Table 1**). We classified FTE, p53 signatures, STIL, and STIC lesions based on consensus pathology criteria that combine H&E histology with p53 and Ki67 immunostaining ^9^. Validation cohorts included an RNA-seq dataset of fimbriae macrodissected from FFPE slides of non-malignant fallopian tubes from 72 women, including 24 premenopausal and 48 postmenopausal (GSE297643, UCLA IRB#20-000551), and a single-cell RNA-seq dataset of 8 premenopausal and 9 postmenopausal non-malignant fallopian tubes (GSE298475, UCLA IRB#22-000091). For computational pathology analysis of fibroblast organization, the validation cohort consisted of 54 digitized H&E-stained fallopian tube slides containing STIC lesionsand adjacent FTE (Cedars-Sinai Medical Center IRB Pro00046215).

### Visium HD data analysis

Visium HD spatial transcriptomic analysis was performed by the UCLA Technology Center for Genomics & Bioinformatics (TCGB). FFPE tissue sections were processed using the 10x Genomics Visium HD Spatial Gene Expression platform following the manufacturer’s protocol ^23^. After fixation and deparaffinization, H&E staining and brightfield imaging were performed. Slides were then destained and decrosslinked before hybridization with a whole transcriptome probe panel. Subsequent steps included probe ligation, CytAssist-enabled probe release, extension, and elution. Final libraries were amplified based on cycle numbers determined by quantitative PCR (qPCR). Library quality and concentration were assessed using an Agilent TapeStation and Qubit fluorometer, respectively, before sequencing on the Illumina NovaSeq X+ platform (paired-end 2×50 bp). Sequencing data were de-multiplexed using BCLConvert v4.3.13 (Illumina). High-resolution (40×) brightfield H&E images were manually aligned to the CytAssist slide image using Loupe Browser (10x Genomics).

### Visium HD dataset preprocessing

FASTQ reads from Visium HD probe-based assay v1 were aligned and quantified using the Space Ranger count pipeline v4.0.1 (10x Genomics) with default settings. We used the Visium Human Transcriptome Probe Set v2.0 with a total of 18085 targeted genes from the GRCh38-2020-A transcriptome. Images were manually aligned via matched CytAssist images in Loupe (v9.0). This edition of Space Ranger provides counts for Visium HD’s 2 µm x 2 µm bins and enables nucleus and cell segmentation based on H&E images using 10x Genomics’ internal StarDist v6 model. Due to varying sample quality, we elected to filter cells and genes for analysis according to the following metrics: greater than 50 unique molecular identifiers (UMIs) per cell, less than 20% of expressed genes were mitochondrial, and genes must be expressed in more than 100 cells. A concatenated SpatialData (v0.5.0) object was created with an AnnData (v0.12.2) table encoding information for 2.7 million cells across 21 samples.

### Cell type annotation

To annotate cell types, we took a hybrid approach that combined unsupervised integration and clustering methods with pathologist-annotated labels. We integrated data from all samples using Harmony-corrected principal components (PCs) ^68^ identified from 8000 highly variable genes (HVGs, scanpy v1.11.4). To reinforce both expressional and spatial relationships in downstream graph clustering, a per-sample joint connectivity nearest-neighbor graph was calculated via a weighted sum of both Harmony-corrected PCs and spatial coordinates. The spatial coordinate connectivity graph is rescaled by the median edge weight of the Harmony-corrected PC connectivity graph before calculating the joint connectivity graph. Using the joint connectivity graph, we clustered cells with the Leiden algorithm at a resolution of 1.5. Resulting clusters were sorted into broad cell types by expression of marker genes.

Pathologist-annotated labels were curated from Space Ranger v4.0.1 graph cluster outputs on a per-sample level. These annotations included graph clusters labels for FTE, STICs, HGSC, STILs, and p53 signatures. However, due to a low presence of STILs and p53 signatures, analysis efforts were focused on comparisons of FTE, STICs, and HGSC. Cells adopted pathologist-annotated labels of FTE, STICs, or HGSC if they were labeled as secretory epithelial cells in unsupervised broad cell type annotations.

DBSCAN, a density-aware clustering algorithm (scikit-learn v1.7.2), was used to identify continuous lesions of FTE, STIC, and HGSC cells. Lesions of STIC and HGSC were clustered together if cells were within 75 µm of one another. Identified lesions were kept if they had a minimum of 30 cells. Lesions of FTE were clustered within 90 µm and a minimum lesion size of 15 cells, due to the thinner epithelial layers of the FTE. Non-lesion FTE, STIC, or HGSC cells were moved to the Unknown category to prevent contamination of spatial analysis.

Subsequent graph-based subclustering with the Louvain method refined cell type annotations. For each subset of cells, we identified 2000 HVGs (Seurat_v3, scanpy v1.11.4), scaled gene expression of log-normalized counts, and identified 50 PCs. These were integrated with Harmony and used to generate a connectivity graph for downstream graph clustering using the Louvain algorithm. Subclustering of secretory epithelial cells, ciliated epithelial cells, and FTE identified ambiguous cell types, most often characterized by incorrect cell segmentation. After pathology-assisted review of ambiguous epithelial cell type subclusters, cells deemed unsuitable for analysis were reclassified as Unknown. Broad immune cell types (T cells, NK cells, B cells, plasma cells, neutrophils, monocytes, macrophages, and mast cells) were similarly identified using this subclustering method.

Immune subtypes, including CD4^+^ T cells, CD8^+^ T cells, M1 macrophages, and M2 macrophages, were annotated using xCell cell type signatures ^69^ and overall expression (OE) gene set scoring. Cell type signatures for CD4^+^ T cells, CD8^+^ T cells, M1 macrophages, and M2 macrophages were created by combining xCell gene set signatures and removing duplicate or absent genes. OE scores are calculated for all cells, and a two-component Gaussian mixture model fits the distribution of cell type signature OE scores within their corresponding broad cell types (T cell or macrophage). These Gaussian components were interpreted as negative and positive expression of the signature. Cells OE scores falling into the positive component were then assigned a posterior probability. For mutually exclusive annotation pairs, such as CD4⁺/CD8⁺ T cells and M1/M2 macrophages, only one identity was retained per cell, selecting the identity with the higher posterior probability.

### Spatial density metric

Density metrics were calculated from the mean distance of 5 nearest-neighbors of type B 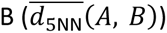 from a center cell A within an interaction radius *r*, generally 100 µm (Eq. 1). These metrics, calculated for each cell across all cell types in the dataset, determined whether cells were considered cell-type-dense. A cell was considered cell-type-dense if its density metric was≥ 0.5, and cell-type-sparse if its density metric was < 0.5. In instances of comparative density analysis, such as comparisons of STIC-dense and FTE-dense cells, a cell must be exclusively STIC-dense or FTE-dense to be included in the analysis.

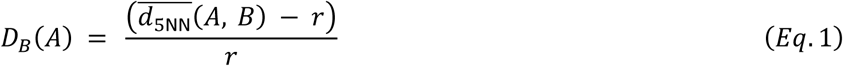

### Differential gene expression analysis

Differentially-expressed genes (DEGs) were identified with the PyDESeq2 algorithm (pydeseq2 v0.5.2). Raw counts for each group were pseudobulked by sum. Sample origin was considered as a covariate to account for batch effects. To prevent incorrect model fitting across cells with different expression patterns, models were only fit on cells included in analysis (Ex. STIC and FTE for STIC versus FTE differential gene expression). A Wald test was then performed to identify DEGs. We then removed a set of ciliated epithelial marker genes, identified from scRNA-seq data of ciliated epithelial cells, and conducted gene set enrichment analysis (GSEA, gseapy v1.1.10) using the following gene sets from MSigDB: Hallmark ^70^ (v2026.1) and Gene Ontology Biological Process ^71^ (v2025.1).

### Cell type composition analysis

Cell type composition of FTE or STIC microenvironments were calculated as a per-sample mean for all cells within a 100 µm radius of FTE and STIC cells, respectively. The 100 µm radius captures biologically plausible local paracrine and immune-mediated interactions. Microenvironment cell composition was also calculated across cells from all samples within a 100 µm radius of FTE and STIC cells. Immune cell composition analysis was calculated as the cross-sample mean of only immune cells within a 100 µm radius of FTE and STIC cells. Cell type proportions are compared at a per-sample level with the Mann-Whitney U test (scipy v1.16.2). P-values are adjusted with the Benjamini-Hochberg (BH) method (statsmodels v0.14.5).

### Stromal cell type composition analysis

Stromal cells contained lower-quality transcriptomic signals that could not always be classified into cell types on a single-cell level. Therefore, stromal cell composition was estimated by pseudobulking unlabeled FTE-, STIC-, and HGSC-associated stroma using CIBERSORTx^39^ with a reference single-cell RNA-seq dataset of stromal cells in healthy human fallopian tubes. Gene expression counts were pseudobulked by sample and spatial association category to create mixture profiles, which were deconvolved by the online version of CIBERSORTx. The resulting cell type proportions were used to determine the overall cell type proportions of the stromal compartment, enabling the comparison of cell compositions between FTE-and STIC-associated stroma. To preserve stromal cells for this analysis, quality control metrics (see Visium HD dataset preprocessing methods) were relaxed by removing the minimum amount of UMIs required. This retained 92.1% of cells.

Epithelial associations to FTE, STIC, or HGSC were assigned using an aforementioned spatial density metric, where association requires classification as an FTE-, STIC-, or HGSC-dense cell within an interaction radius of 200 µm (see spatial density metric methods). Cells are restricted to a single association based on the highest density score. The CIBERSORTx mixture file contained FTE- and STIC-associated stromal cells pseudobulked into separate columns, with gene expression representing the rows. The mixtures were deconvolved against a signature matrix that was derived from scRNA-seq analysis of stromal cells in healthy human fallopian tubes ^72^. Both the mixture file and signature matrix were normalized to counts per million prior to analysis. CIBERSORTx was run with 1000 permutations using bulk batch corrections without a source GEP file because quantile normalization was disabled. CIBERSORTx outputs of reference cell type proportions were mapped to Visium HD annotations by matching transcriptomically similar cell types before integration for cell type proportion analysis.

### Radial enrichment analysis

Radial enrichment plots are calculated based on lesion annotations of FTE, STIC, and HGSC. We find the proportion of all cells within 100 µm of FTE cells based on their distance from the FTE reference cell, and bin them into 50 bins of radius 2 µm. To prevent division by zero, Laplace smoothing was applied (Eq. 2) with *α* = 0.0001. We performed this analysis on bins from 10-100 µm, ignoring bins that are potentially smaller than the diameter of a cell (∼10 µm).

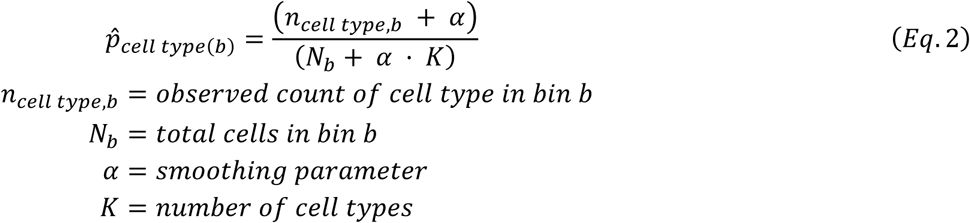

On a per-sample basis, we treat STIC and HGSC cells as alternative anchor cells and compare them to the baseline neighborhood of FTE (Eq. 3). Enrichment greater than 0 on a log 2 scale indicates that a given cell type is more common in the anchor cell microenvironment than FTE microenvironments at that distance, and vice versa.

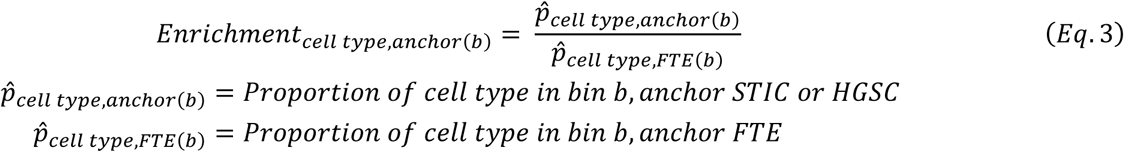

### Spatial niche analysis

Unique niches, or microenvironments, are identified through k-means clustering (scikit-learn v1.7.2) of density metrics. Broad microenvironments were identified from all cells in the dataset (**Supp. Fig. 8A**). **Supp. Fig. 8B** plots z-score normalized median density for each niche, normalized across a given cell type density. Epithelial enrichment values are calculated as the niche FTE and STIC proportions divided by the observed proportion of FTE and STIC across the included dataset. Cell counts of FTE and STIC are plotted in a butterfly plot with diamonds indicating the expected cell counts for each niche, given the observed proportion of FTE and STIC.

Epithelial niches were identified in a similar manner. However, FTE and STIC labels were grouped into one epithelial category to find microenvironments that were shared or differed between each microenvironment. Only cells that were STIC-or FTE-dense were included in this analysis. Epithelial enrichment values were calculated as the niche FTE and STIC proportions divided by the observed proportion of FTE and STIC within the selected STIC- or FTE-dense cells. Similarly, expected cell counts are based on the observed proportion of FTE and STIC within the selected population of STIC- or FTE-dense cells.

### Ligand receptor analysis

Ligand receptor analysis consisted of two spatially-subsetted permutation tests, hereafter referred to as epithelial static and target cell static. Target cell types are classified as broad labels of T cells, macrophages, and fibroblasts. These tests respectively seek to answer two questions: 1) Do FTE/STIC cells that are proximal to target cells (T cells, macrophages, and fibroblasts, **Suppl. Fig. 8C**) express unique signaling patterns, altering the microenvironment?, and 2) do target cells that are proximal to FTE/STIC cells interact differently compared to other epithelial cells, altering epithelial response?

In the epithelial static test, proximal target cells are defined as target cells with a respective FTE/STIC density metric ≥ 0.5. Proximal FTE/STIC cells are defined as FTE/STIC cells with a respective target cell density metric ≥ 0.5 and FTE/STIC density metric ≥ 0.5. We then hold target cell-proximal FTE/STIC cells static while shuffling all other cell type labels, measuring the randomized mean co-expression of FTE/STIC-proximal target cells and target cell-proximal FTE/STIC cells 1000 times. This creates a randomized null distribution of mean co-expression, against which we can test the initially observed mean co-expression to identify our p-values. P-values are adjusted with the Benjamini-Hochberg method.

For the target cell static test, we held FTE/STIC-proximal target cells static and shuffled all other epithelial labels (Ciliated, FTE, STIC, HGSC, and Unknown). Unknown cells contained cells that did not meet the pathologist’s definition of an epithelial cell type label or discontinuous epithelial labels. However, their transcriptomic profile was similar to that of other epithelial cells in the dataset. Similar to the epithelial static test, we created a null distribution of mean co-expression of FTE/STIC-proximal target cells and target cell-proximal FTE/STIC cells by permuting cell type labels 1000 times. We then tested our initially observed mean co-expression against the null distribution to identify our p-values. P-values are adjusted with the BH method (statsmodels v0.14.5).

Permutation tests were performed on a per-sample level to maintain spatial relevance. Results were aggregated across samples, grouped by test type. Ligand-receptor pairs were identified from CellChatDB v2 ^73^, a curated database of ligand-receptor pairs containing experimentally validated and annotated ligand-receptor pairs, as well as the pathways these pairs are associated with. Interactions that were statistically significant in more than 4 samples were kept for downstream analysis (n=8). For heteromeric complexes, where multiple receptors are required for cell-cell interactions, we initially split these interactions into individual ligand-receptor pairs. Then, when summarizing the significant pathways, heteromeric complexes are only included if their split ligand-receptor pairs were both significant.

### Overall Expression (OE) gene set scoring

Gene signature activity was scored on a per-cell basis using an Overall Expression algorithm adapted from Jerby-Arnon et al. ^74^, designed to control for expression-level bias in sparse single-cell data. For each sample, all genes are ranked into 40 bins by mean expression, such that genes within a bin have comparable baseline expression levels.

For a cell c and respective gene set signature S, the observed score is:

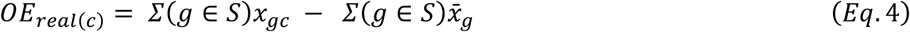

where: *x_gc_* is the expression of gene *g* in cell *c* and *x̄*_*g*_ is the mean expression of gene g across the sample. To compute the expected score, the bin composition of *S* is matched and random gene sets of identical bin composition are drawn 500 times:

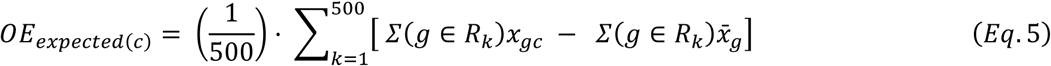

where: *R_k_* is the *k*-th random gene set. The final score is:

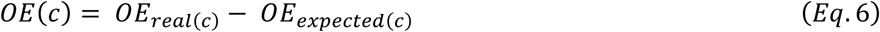

An OE pathway score > 0 indicates that the cell expresses the signature above random expectation. Less than 0 indicates below, and 0 is equivalent to random.

Gene set pathway analysis for each cell utilized select Hallmark gene sets from the MSigDB Hallmark gene set collection (v2026.1). Median OE pathway scores for each pathway were compared between all cell types considered STIC-dense or FTE-dense with a Mann-Whitney U test (scipy v1.16.2) on per-sample median OE scores. P-values are adjusted with the BH method (statsmodels v0.14.5). Cell type-sample combinations of at least 3 unique samples with at least 5 cells in each cell group were included. Additionally, at least 20 total cells were required for analysis.

To further assess the relationship between cell type-specific pathway activity and FTE-, STIC-, or HGSC-density, we fit linear mixed-effects models using the MixedLM function from statsmodels (v0.14.5) between all cell types and OE pathway scores. OE pathway scores were modeled as the response variable with density as a fixed-effect predictor and sample as a random effect to account for inter-sample variability. Only cell type-sample combinations with at least 3 unique samples and 20 total cells were included. Fixed-effect coefficients and p-values were extracted from each model fit, and p-values were corrected for multiple comparisons with the BH method (statsmodels v0.14.5).

### Bulk RNA sequencing

FFPE blocks from 74 specimens (normal fallopian tubes) were sectioned onto uncharged glass slides. One 4 μm H&E-stained section was used by a pathologist to circle the fimbriated area of the fallopian tube. Depending on the size, 1–3 unstained 10 μm sections were macrodissected with a clean razor blade. Total RNA was isolated using the miRNeasy FFPE kit according to the manufacturer’s instructions (Qiagen). Library prep was performed using the SMARTer Stranded Total RNA-Seq Kit v2. Unwanted sequences (non-polyA-tailed RNAs from the sample, mitochondrial genome sequences, ribosomal RNAs, transfer RNAs, adapter sequences and others) were removed using Bowtie2 (version 2.2.4). The paired-end reads were aligned to the reference human genome downloaded from the UCSC database (GRCh37/hg19). STAR (2.4.1) aligner was used for read alignment. Reads mapping to ribosomal and mitochondrial genomes were removed before alignment was performed. The raw read counts were estimated using HTSeq-0.6.1. Data normalization and differential expression were done using DESeq2.

### Single-cell RNA sequencing

Fallopian tubes from 17 patients were collected by the UCLA Pathology team, placed in DMEM media, and transferred on ice to the lab for further processing. Preparation for single-cell RNA sequencing was done as previously described ^75^. Partek flow software was used for QC, thresholding, and data analysis. Cells were removed according to the following criteria: 1) cells had fewer than 500 genes or more than 10000 genes; 2) cells had fewer than 500 unique molecular identifiers (UMI) or over 10000 UMI; and 3) cells had more than 10% of mitochondrial UMI counts. For cell identification, we used the following markers: Epithelial cells: EPCAM, KRT18, PAX8, FOXJ1; Fibroblast: PDGFRA, COL1A1; T and NK cells: CD8A, CD3E, NKG7; Macrophages: C1QA, CD68; Mast cells: TPSAB1; Endothelial Cells: CLDN5, CDH5; B Cells: JCHAIN, CD79A; Neutrophils: S1008A, S100A9. Epithelial cells were re-clustered to identify ciliated and secretory cell clusters.

### Immunohistochemistry and antibodies

Immunohistochemistry on formalin-fixed paraffin-embedded tissue sections was performed by the UCLA Translational Pathology Core Laboratory (TPCL) using the Leica Bond RX processor based on Protocol F using Bond Polymer Refine Detection kit (Cat#DS9800, Leica Biosystems, Nussloch, Germany), ER2 buffer (BOND Epitope Retrieval Solution 2, Leica Biosystems, Cat#AR9640), DakoCytomation Envision System Labelled Polymer HRP anti rabbit (Agilent, Santa Clara, CA, USA, K4003) and antibodies against MUC6 (AB223846, Abcam), ASF1B (HPA069385, Sigma), PRAME (NBP3-45107, Novus Biologicals), FOXM1 (WH0002305M2, Sigma), TPX2 (HPA005487, Sigma), S100A4 (F4023, Selleckchem), EZH2 (F0281, Selleckchem), PLK1 (F0393, Selleckchem), UCHL1 (13179, Cell Signaling), SCGB1A1 (10490-1-AP, Proteintech), MGP (MA5-26793, Invitrogen), and SYNE1 (HPA019113, Sigma). Hematoxylin was used as the counterstain.

### ASF1B knockout generation and in vitro cell line analyses

SKOV3ip1 control and ASF1B knockout (KO) cell lines were generated using a lentiviral vector co-expressing wild-type SpCas9 and an sgRNA targeting human ASF1B (Entrez ID: 55723; exon 1 target sequence: 5′-ACTCGAAGCTGATCTCGAAC-3′) obtained from Horizon Discovery (Edit-R All-in-One Lentiviral sgRNA system). The vector was transfected into cells, and 2 days later, the medium was replaced with fresh medium containing 1 to 2 μg/mL puromycin for selection of stably transfected cells. Single-cell clones were isolated and screened for ASF1B loss by western blotting using an anti-ASF1B antibody, and by qPCR using specific primers: Forward: 3’TCCGGTTCGAGATCAGCTTC 5’;Reverse: 3’GTCGGCCTGAAAGACAAACA 5’. We assessed whether ASF1B KO affected ASF1A expression by qPCR using especific primers: Forward: 3’TGCTGGATAACCCTTCTCCTT 5’; Reverse: 3’CAGGACCCACTAAAACAGAGTC 5’ Western blotting, qPCR, cell proliferation, and migration assays were performed as previously described ^76, 77^.

### Subcutaneous tumor growth

All mouse procedures were conducted in accordance with the NIH Guide for the Care and Use of Laboratory Animals and were approved by the University of California, Los Angeles (UCLA) Animal Research Committee (ARC). Eight-week-old athymic nude (Nu/Nu) mice were purchased from Charles River (Strain #088). For tumor growth experiments, 1 × 10^6^ SKOV3ip1 control or ASF1B KO cells were injected subcutaneously into the bilateral flanks of 10 mice per group (N=20). Tumor growth was monitored for 40 days, until the first signs of ulceration appeared in the control group. Tumor volume was measured with calipers twice weekly throughout the experiment.

### Computational pathology approach to quantify collagen fiber organization

H&E slides were scanned at 40× magnification using an Aperio AT Turbo slide scanner (Leica Biosystems). A pathologist annotated the digitized H&E slides using ImageScope software (Leica Biosystems). In the discovery cohort (n = 9 slides with STIC lesions from the Visium HD cohort), stromal regions adjacent to STIC lesions and FTE were annotated on each slide. In the validation cohort (n = 54 H&E slides with malignant lesions in the fallopian acquired at the Cedars-Sinai Medical Center in Los Angeles), stromal regions adjacent to HGSC, STIC, and FTE were annotated, with one to three regions of each type per slide when present. Unlike the discovery cohort, in which each slide contained stromal regions adjacent to both STIC and FTE, the validation cohort consisted of slides containing at least one stromal region adjacent to HGSC, STIC, or FTE, not necessarily all three on the same slide.

Annotated stromal regions were sampled using 512 × 512-pixel regions of interest (ROIs) for downstream quantification of collagen fiber organization. Each ROI underwent color deconvolution ^78^ to isolate a grayscale image corresponding to eosinophilic staining for downstream analysis. Grayscale images were processed using derivative-of-Gaussian filters that we previously used ^79, 80^ to generate fiber masks that detect and delineate collagen fibers. Collagen organization was quantified using four features extracted from the fiber masks: median fiber length, standard deviation of fiber length, fiber density (defined as the ratio of fiber pixels to total pixels within the ROI), and fiber orientation entropy ^45^. For each case, feature values were summarized as the median across ROIs of the same tissue type within a slide. Before feature extraction, H&E staining variability across ROIs was normalized using Reinhard’s color standardization method ^81^.

### Statistical analysis of collagen fiber organization

We first performed a case-level analysis (discovery set) using paired Wilcoxon signed-rank tests across the 9 paired median values (STIC vs FTE) for each feature. For each comparison, we report p-values, median differences, and paired effect sizes estimated using Cohen’s *d*. To account for multiple testing across the four features, p-values were adjusted using the Benjamini–Hochberg false discovery rate controlling procedure with a threshold of 0.05. Effect sizes were visualized using a forest plot showing per-case differences (median(STIC) − median(FTE)) derived from ROI-level data. Ninety-five percent confidence intervals were estimated by paired bootstrap resampling of medians (2,000 iterations). Using the bootstrapped confidence intervals and per-case effect estimates, the pooled heterogeneity statistic (I² - using the DerSimonian-Laird method of moments) was calculated to quantify the proportion of total variability in effect sizes attributable to true between-case differences rather than sampling error.

Significant features that differed between STIC-associated stroma and FTE-associated stroma in the discovery set were validated in the validation set using a linear mixed-effects model. The model was defined as: feature ∼ ROI_type + (1 | Case), where: ROI_type (FTE, STIC, HGSC) was included as a fixed effect to evaluate differences across ROIs, and Case was modeled as a random intercept to account for H&E slide-level variability. FTE-associated stroma was used as the reference category, yielding fixed-effect estimates for the STIC–FTE and HGSC–FTE contrasts, and an additional linear contrast for STIC–HGSC. Variance components (between-case variance: σ²_case_, and within-case ROI-level variance: σ²_ROI_) were used to compute the intraclass correlation coefficient (ICC), quantifying the proportion of variance attributable to case-level differences. Model fit for the linear mixed-effects analyses was assessed using the Akaike information criterion (AIC). As a secondary, exploratory analysis, per-case median feature values were computed, and differences between ROI categories were evaluated using the Mann–Whitney U test. Features reaching significance were visualized across FTE, STIC, and HGSC groups.

## Data availability

Datasets generated during the current study are available in the GEO database: <u>GSE303022</u>– Visium HD dataset of 21 capture areas featuring areas of normal fallopian tubes, STIC lesions and invasive ovarian carcinoma GSE297643 - Bulk RNA-seq dataset of 74 normal premenopausal and postmenopausal fallopian tubes.

GSE298475- Single Cell RNA-seq dataset of 17 normal premenopausal and postmenopausal fallopian tubes.

## CRediT authorship contribution statement

**MSR:** Writing – original draft, Methodology, Formal analysis, Visualization, Validation, Data curation, Conceptualization.

**KT:** Writing – original draft, Methodology, Formal analysis, Visualization, Validation, Data curation, Conceptualization.

**AS:** Writing – review & editing, Investigation, Formal analysis, Visualization, Validation, Data curation.

**SK:** Writing – review & editing, Resources, Methodology.

**DP:** Writing – review & editing, Methodology, Formal analysis.

**YR:** Writing – review & editing, Investigation, Methodology, Formal analysis, Data curation.

**BTH:** Writing – review & editing, Methodology, Investigation.

**JHK:** Writing – review & editing, Methodology, Formal analysis.

**RM:** Writing – review & editing, Methodology, Formal analysis.

**MTR:** Writing – review & editing, Methodology, Formal analysis.

**RF:** Writing – review & editing, Methodology.

**DL:** Writing – review & editing, Methodology, Formal analysis, Software resources.

**EJF:** Writing – review & editing, Methodology, Software resources.

**AEW:** Writing – review & editing, Resources, Data curation.

**AG:** Writing – original draft, Software resources, Methodology, Formal analysis, Visualization, Data curation.

**BYK:** Writing – review & editing, Resources, Data curation, Conceptualization.

**AMX:** Writing – original draft, Software resources, Visualization, Validation, Supervision, Project administration, Methodology, Investigation, Formal analysis, Data curation, Conceptualization.

**SO:** Writing – original draft, Visualization, Validation, Supervision, Resources, Project administration, Methodology, Investigation, Formal analysis, Data curation, Conceptualization.

## Acknowledgements

We thank the UCLA cores TPCL and TCGB for their assistance with this research. We are deeply grateful to the women who had the foresight and generosity to consent to the use of their clinical samples for research.

## REFERENCES

1. Siegel RL, Kratzer TB, Wagle NS, Sung H, Jemal A. Cancer statistics, 2026. CA Cancer J Clin. 2026;76: e70043.

2. Balkwill FR, Laumont CM, Burdett N, et al. Rethinking ovarian cancer III: the past decade and future directions. Nat Rev Cancer. 2026.

3. Kurman RJ, Shih I-M. Molecular pathogenesis and extraovarian origin of epithelial ovarian cancer--shifting the paradigm. Hum Pathol. 2011;42: 918–931.

4. Labidi-Galy SI, Papp E, Hallberg D, et al. High grade serous ovarian carcinomas originate in the fallopian tube. Nat Commun. 2017;8: 1093.

5. Recouvreux MS, Orsulic S. Before They Were Malignant. Cancer Discov. 2025;15: 1093–1095.

6. Powell CB, Kenley E, Chen L-m, et al. Risk-reducing salpingo-oophorectomy in BRCA mutation carriers: role of serial sectioning in the detection of occult malignancy. Journal of Clinical Oncology. 2005;23: 127–132.

7. Medeiros F, Muto MG, Lee Y, et al. The tubal fimbria is a preferred site for early adenocarcinoma in women with familial ovarian cancer syndrome. American Journal of Surgical Pathology. 2006;30: 230–236.

8. Vang R, Visvanathan K, Gross A, et al. Validation of an algorithm for the diagnosis of serous tubal intraepithelial carcinoma. International Journal of Gynecological Pathology. 2012;31: 243–253.

9. Bogaerts JMA, van Bommel MHD, Hermens R, et al. Consensus based recommendations for the diagnosis of serous tubal intraepithelial carcinoma: an international Delphi study. Histopathology. 2023;83: 67–79.

10. Kuhn E, Kurman RJ, Vang R, et al. TP53 mutations in serous tubal intraepithelial carcinoma and concurrent pelvic high-grade serous carcinoma-evidence supporting the clonal relationship of the two lesions. Journal of Pathology. 2012;226: 421–426.

11. Kindelberger DW, Lee Y, Miron A, et al. Intraepithelial carcinoma of the fimbria and pelvic serous carcinoma: evidence for a causal relationship. American Journal of Surgical Pathology. 2007;31: 161–169.

12. Zhang S, Dolgalev I, Zhang T, Ran H, Levine DA, Neel BG. Both fallopian tube and ovarian surface epithelium are cells-of-origin for high-grade serous ovarian carcinoma. Nat Commun. 2019;10: 5367.

13. Akahane T, Masuda K, Hirasawa A, et al. TP53 variants in p53 signatures and the clonality of STICs in RRSO samples. Journal of Gynecologic Oncology. 2022;33: e50.

14. Kader T, Lin JR, Hug CB, et al. Multimodal Spatial Profiling Reveals Immune Suppression and Microenvironment Remodeling in Fallopian Tube Precursors to High-Grade Serous Ovarian Carcinoma. Cancer Discov. 2025;15: 1180–1202.

15. Garcia GL, Orellana T, Gorecki G, et al. Aged and BRCA mutated stromal cells drive epithelial cell transformation. Cancer Discov. 2025.

16. Wang Y, Huang P, Wang BG, et al. Spatial Transcriptomic Analysis of Ovarian Cancer Precursors Reveals Reactivation of IGFBP2 during Pathogenesis. Cancer Res. 2022;82: 4528–4541.

17. Chang TY, Chien YW, Chen SH, et al. Integrated Spatial Analysis Reveals the Molecular Landscape of Ovarian Precancerous Lesions. Cancer Res. 2026;86: 1739–1752.

18. Wang Y, Douville C, Chien YW, et al. Aneuploidy Landscape in Precursors of Ovarian Cancer. Clin Cancer Res. 2024;30: 600–615.

19. Makhmut A, Dragomir MP, Fritzsche S, et al. Spatial proteomics of ovarian cancer precursors delineates early disease changes and drug targets. Mol Syst Biol. 2026;22: 7–41.

20. Janesick A, Shelansky R, Gottscho AD, et al. High resolution mapping of the tumor microenvironment using integrated single-cell, spatial and in situ analysis. Nat Commun. 2023;14: 8353.

21. He S, Bhatt R, Brown C, et al. High-plex imaging of RNA and proteins at subcellular resolution in fixed tissue by spatial molecular imaging. Nat Biotechnol. 2022;40: 1794–1806.

22. Ståhl PL, Salmén F, Vickovic S, et al. Visualization and analysis of gene expression in tissue sections by spatial transcriptomics. Science. 2016;353: 78–82.

23. Oliveira MF, Romero JP, Chung M, et al. High-definition spatial transcriptomic profiling of immune cell populations in colorectal cancer. Nat Genet. 2025;57: 1512–1523.

24. Balakrishnan K, Chen Y, Dong J. Amplified Cell Cycle Genes Identified in High-Grade Serous Ovarian Cancer. Cancers (Basel). 2024;16.

25. Kobayashi H, Ogawa K, Kawahara N, et al. Sequential molecular changes and dynamic oxidative stress in high-grade serous ovarian carcinogenesis. Free Radic Res. 2017;51: 755–764.

26. Bijron JG, Seldenrijk CA, Zweemer RP, Lange JG, Verheijen RH, van Diest PJ. Fallopian tube intraluminal tumor spread from noninvasive precursor lesions: a novel metastatic route in early pelvic carcinogenesis. Am J Surg Pathol. 2013;37: 1123–1130.

27. Colina JA, Recouvreux MS, Sobeck AM, et al. Ciliated Cells Drive Critical STING-Mediated Tumor Suppression in the Fallopian Tube Epithelium. Cancer Res. 2026: Of1-of18.

28. Siraj Y, Aprile D, Alessio N, Peluso G, Di Bernardo G, Galderisi U. IGFBP7 is a key component of the senescence-associated secretory phenotype (SASP) that induces senescence in healthy cells by modulating the insulin, IGF, and activin A pathways. Cell Commun Signal. 2024;22: 540.

29. Al Shboul S, Awad H, Abu-Humaidan A, Ababneh NA, Khasawneh AI, Saleh T. Oncogene-Induced Senescence Transcriptomes Signify Premalignant Colorectal Adenomas. Curr Issues Mol Biol. 2025;47.

30. Zheng D, Limmon GV, Yin L, et al. Regeneration of alveolar type I and II cells from Scgb1a1-expressing cells following severe pulmonary damage induced by bleomycin and influenza. PLoS One. 2012;7: e48451.

31. Rawlins EL, Okubo T, Xue Y, et al. The role of Scgb1a1+ Clara cells in the long-term maintenance and repair of lung airway, but not alveolar, epithelium. Cell Stem Cell. 2009;4: 525–534.

32. Zheng D, Yin L, Chen J. Evidence for Scgb1a1(+) cells in the generation of p63(+) cells in the damaged lung parenchyma. Am J Respir Cell Mol Biol. 2014;50: 595–604.

33. Khatri A, Kraft BD, Tata PR, Randell SH, Piantadosi CA, Pendergast AM. ABL kinase inhibition promotes lung regeneration through expansion of an SCGB1A1+ SPC+ cell population following bacterial pneumonia. Proc Natl Acad Sci U S A. 2019;116: 1603–1612.

34. Kim CF, Jackson EL, Woolfenden AE, et al. Identification of bronchioalveolar stem cells in normal lung and lung cancer. Cell. 2005;121: 823–835.

35. Song H, Weinstein HNW, Allegakoen P, et al. Single-cell analysis of human primary prostate cancer reveals the heterogeneity of tumor-associated epithelial cell states. Nat Commun. 2022;13: 141.

36. Huang FW, Song H, Weinstein HN, et al. Club-like cells in proliferative inflammatory atrophy of the prostate. J Pathol. 2023;261: 85–95.

37. Quintar AA, Mukdsi JH, del Valle Bonaterra M, Aoki A, Maldonado CA, Pérez Alzaa J. Increased expression of uteroglobin associated with tubal inflammation and ectopic pregnancy. Fertil Steril. 2008;89: 1613–1617.

38. Arai J, Hayakawa Y, Tateno H, Fujiwara H, Kasuga M, Fujishiro M. The role of gastric mucins and mucin-related glycans in gastric cancers. Cancer Sci. 2024;115: 2853–2861.

39. Steen CB, Liu CL, Alizadeh AA, Newman AM. Profiling Cell Type Abundance and Expression in Bulk Tissues with CIBERSORTx. Methods Mol Biol. 2020;2117: 135–157.

40. Metousis A, Kenny HA, Shimizu A, et al. Integration of cell-type resolved spatial proteomics and transcriptomics reveals novel mechanisms in early ovarian cancer. medRxiv. 2025.

41. Newton JB, Weiss SN, Nuss CA, et al. Decorin and/or biglycan knockdown in aged mouse patellar tendon impacts fibril morphology, scar area, and mechanical properties. J Orthop Res. 2024;42: 2400–2413.

42. Chen D, Smith LR, Khandekar G, et al. Distinct effects of different matrix proteoglycans on collagen fibrillogenesis and cell-mediated collagen reorganization. Sci Rep. 2020;10: 19065.

43. Wan Z, Bai X, Wang X, et al. Mgp High-Expressing MSCs Orchestrate the Osteoimmune Microenvironment of Collagen/Nanohydroxyapatite-Mediated Bone Regeneration. Adv Sci (Weinh). 2024;11: e2308986.

44. Wu J, Raz Y, Recouvreux MS, et al. Focal Serous Tubal Intra-Epithelial Carcinoma Lesions Are Associated With Global Changes in the Fallopian Tube Epithelia and Stroma. Front Oncol. 2022;12: 853755.

45. Li H, Bera K, Toro P, et al. Collagen fiber orientation disorder from H&E images is prognostic for early stage breast cancer: clinical trial validation. NPJ Breast Cancer. 2021;7: 104.

46. Acland M, Arentz G, Mussared M, et al. Proteomic Analysis of Pre-Invasive Serous Lesions of the Endometrium and Fallopian Tube Reveals Their Metastatic Potential. Front Oncol. 2020;Volume 10 - 2020.

47. da Silva EM, Da Cruz Paula A, Dessources K, et al. A Subset of Serous Tubal Intraepithelial Carcinoma (STIC)-Like Lesions and Concurrent High-Grade Endometrial Carcinoma Are Genomically Related Entities. Mod Pathol. 2025;38: 100890.

48. Emori MM, Drapkin R. The hormonal composition of follicular fluid and its implications for ovarian cancer pathogenesis. Reprod Biol Endocrinol. 2014;12: 60.

49. Arai J, Hayakawa Y, Tateno H, et al. Impaired Glycosylation of Gastric Mucins Drives Gastric Tumorigenesis and Serves as a Novel Therapeutic Target. Gastroenterology. 2024;167: 505–521.e519.

50. Zheng H, Takahashi H, Nakajima T, et al. MUC6 down-regulation correlates with gastric carcinoma progression and a poor prognosis: an immunohistochemical study with tissue microarrays. J Cancer Res Clin Oncol. 2006;132: 817–823.

51. Hung CH, Chen LC, Zhang Z, et al. Regulation of TH2 responses by the pulmonary Clara cell secretory 10-kd protein. J Allergy Clin Immunol. 2004;114: 664–670.

52. Pilon AL. Rationale for the development of recombinant human CC10 as a therapeutic for inflammatory and fibrotic disease. Ann N Y Acad Sci. 2000;923: 280–299.

53. Levine CR, Gewolb IH, Allen K, et al. The safety, pharmacokinetics, and anti-inflammatory effects of intratracheal recombinant human Clara cell protein in premature infants with respiratory distress syndrome. Pediatr Res. 2005;58: 15–21.

54. Linz VC, Löwe A, van der Ven J, Hasenburg A, Battista MJ. Incidence of pelvic high-grade serous carcinoma after isolated STIC diagnosis: A systematic review of the literature. Front Oncol. 2022;12: 951292.

55. van den Berg CB, Dasgupta S, Ewing-Graham PC, et al. Does serous tubal intraepithelial carcinoma (STIC) metastasize? The clonal relationship between STIC and subsequent high-grade serous carcinoma in BRCA1/2 mutation carriers several years after risk-reducing salpingo-oophorectomy. Gynecol Oncol. 2024;187: 113–119.

56. Steenbeek MP, van Bommel MHD, Bulten J, et al. Risk of Peritoneal Carcinomatosis After Risk-Reducing Salpingo-Oophorectomy: A Systematic Review and Individual Patient Data Meta-Analysis. J Clin Oncol. 2022;40: 1879–1891.

57. Negri S, Fisch C, de Hullu JA, et al. Diagnosis and management of isolated serous tubal intraepithelial carcinoma: A qualitative focus group study. Bjog. 2024;131: 1851–1861.

58. Konecny GE, Hendrickson AEW, Winterhoff B, et al. 721MO Phase I, two-part, multicenter first-in-human (FIH) study of TORL-1-23: A novel claudin 6 (CLDN6) targeting antibody drug conjugate (ADC) in patient with advanced solid tumors. Annals of Oncology. 2024;35: S551.

59. Kaczorowski M, Chłopek M, Kruczak A, Ryś J, Lasota J, Miettinen M. PRAME Expression in Cancer. A Systematic Immunohistochemical Study of >5800 Epithelial and Nonepithelial Tumors. Am J Surg Pathol. 2022;46: 1467–1476.

60. Mackensen A, Haanen J, Koenecke C, et al. CLDN6-specific CAR-T cells plus amplifying RNA vaccine in relapsed or refractory solid tumors: the phase 1 BNT211-01 trial. Nat Med. 2023;29: 2844–2853.

61. Reinhard K, Rengstl B, Oehm P, et al. An RNA vaccine drives expansion and efficacy of claudin-CAR-T cells against solid tumors. Science. 2020;367: 446–453.

62. Holmberg-Thydén S, Dufva IH, Gang AO, et al. Epigenetic therapy in combination with a multi-epitope cancer vaccine targeting shared tumor antigens for high-risk myelodysplastic syndrome - a phase I clinical trial. Cancer Immunol Immunother. 2022;71: 433–444.

63. Pujol JL, De Pas T, Rittmeyer A, et al. Safety and Immunogenicity of the PRAME Cancer Immunotherapeutic in Patients with Resected Non-Small Cell Lung Cancer: A Phase I Dose Escalation Study. J Thorac Oncol. 2016;11: 2208–2217.

64. Weber JS, Vogelzang NJ, Ernstoff MS, et al. A phase 1 study of a vaccine targeting preferentially expressed antigen in melanoma and prostate-specific membrane antigen in patients with advanced solid tumors. J Immunother. 2011;34: 556–567.

65. Gutzmer R, Rivoltini L, Levchenko E, et al. Safety and immunogenicity of the PRAME cancer immunotherapeutic in metastatic melanoma: results of a phase I dose escalation study. ESMO Open. 2016;1: e000068.

66. Konecny GE, Hendrickson AEW, Winterhoff B, et al. Initial results of dose finding in a first-in-human phase 1 study of a novel Claudin 6 (CLDN6) targeted antibody drug conjugate (ADC) TORL-1-23 in patients with advanced solid tumors. Journal of Clinical Oncology. 2023;41: 3082–3082.

67. Gant KL, Jambor AN, Li Z, et al. Evaluation of Collagen Alterations in Early Precursor Lesions of High Grade Serous Ovarian Cancer by Second Harmonic Generation Microscopy and Mass Spectrometry. Cancers (Basel). 2021;13.

68. Korsunsky I, Millard N, Fan J, et al. Fast, sensitive and accurate integration of single-cell data with Harmony. Nat Methods. 2019;16: 1289–1296.

69. Aran D, Hu Z, Butte AJ. xCell: digitally portraying the tissue cellular heterogeneity landscape. Genome Biol. 2017;18: 220.

70. Liberzon A, Birger C, Thorvaldsdóttir H, Ghandi M, Mesirov JP, Tamayo P. The Molecular Signatures Database (MSigDB) hallmark gene set collection. Cell Syst. 2015;1: 417–425.

71. The Gene Ontology knowledgebase in 2026. Nucleic Acids Res. 2026;54: D1779–d1792.

72. Ulrich ND, Shen YC, Ma Q, et al. Cellular heterogeneity of human fallopian tubes in normal and hydrosalpinx disease states identified using scRNA-seq. Dev Cell. 2022;57: 914–929.e917.

73. Jin S, Plikus MV, Nie Q. CellChat for systematic analysis of cell-cell communication from single-cell transcriptomics. Nat Protoc. 2025;20: 180–219.

74. Jerby-Arnon L, Shah P, Cuoco MS, et al. A Cancer Cell Program Promotes T Cell Exclusion and Resistance to Checkpoint Blockade. Cell. 2018;175: 984–997.e924.

75. Hu Y, Recouvreux MS, Haro M, et al. INHBA(+) cancer-associated fibroblasts generate an immunosuppressive tumor microenvironment in ovarian cancer. NPJ Precis Oncol. 2024;8: 35.

76. Recouvreux MS, Miao J, Gozo MC, et al. FOXC2 Promotes Vasculogenic Mimicry in Ovarian Cancer. Cancers (Basel). 2022;14.

77. Recouvreux MS, Taylor-Harding B, Rowat AC, Karlan BY, Orsulic S. Cancer Growth and Invasion Are Increased in the Tight Skin (TSK) Mouse. Cancers (Basel). 2025;17.

78. Ruifrok AC, Johnston DA. Quantification of histochemical staining by color deconvolution. Anal Quant Cytol Histol. 2001;23: 291–299.

79. Griffin LD, Lillholm M. Symmetry Sensitivities of Derivative-of-Gaussian Filters. IEEE Transactions on Pattern Analysis and Machine Intelligence. 2010;32: 1072–1083.

80. Smick AH, Zurek N, Taylor-Harding B, et al. Molecular and spatial differences in the tumor microenvironment of high-grade serous ovarian cancers with short versus long-term survival. Gynecol Oncol. 2026;210: 55–62.

81. Reinhard E, Adhikhmin M, Gooch B, Shirley P. Color transfer between images. IEEE Computer Graphics and Applications. 2001;21: 34–41.

