## Supplemental Figures for "Spatially resolved transcriptional programs link fallopian tube precursor lesions to immune activation and stromal reorganization"

#### SUPPLEMENTARY TABLE LEGENDS

**Suppl. Table 1.** Patient cohort - Visium HD dataset (GSE303022) (OVCA= ovarian cancer; HRD= homologous recombination deficient; VUS= variant of unknown significance; HRT= hormone replacement therapy).
**Suppl. Table 2.** Curated ciliated cell gene expression
**Suppl. Table 3.** Genes differentially expressed in STIC vs FTE.
**Suppl. Table 4.** STIC vs FTE GSEA.

#### SUPPLEMENTARY FIGURE LEGENDS

**Suppl. Fig. 1. Visium HD identifies 20 unique cell types across 21 fallopian tube samples.**
**A.** Uniform manifold approximation and projection (UMAP) of the concatenated 21-sample Visium HD dataset.
**B.** Cell counts across all 21 samples and 20 cell types.
**C.** Cell counts for each sample after QC.
**D.** Distribution of total UMIs (top), genes (middle), and mitochondrial genes (bottom) in each sample. These metrics were calculated for both the pre-QC (left) and post-QC (right) datasets.

**Suppl. Fig. 2. IHC validation of genes upregulated in STIC lesions.** Thick arrows indicate STIC lesions, whereas thin arrows denote FTE.

**Suppl. Fig. 3. A case containing both a STIL and STIC lesion shows that genes identified as upregulated in STIC relative to FTE are not similarly upregulated in the STIL lesion from the same patient.**
**A-B.** High (A) and low (B) magnification views of an H&E-stained section of a fallopian tube fimbria from the same patient containing a STIL and STIC lesion
**C.** Visium HD spatial transcriptomic overlay of 209 genes identified as upregulated in STIC relative to FTE with log2 fold change > 2.

**Suppl. Fig. 4. ASF1B loss suppresses the tumorigenic properties of SKOV3ip1 human ovarian cancer cells.**
**A.** Western blot analysis of control and ASF1B knockout SKOV3ip1 cells.
**B.** qPCR analysis of ASF1B (left) and ASF1A (right) expression in control and knockout SKOV3ip1 cells. *t*-test, ****$p < 0.001$.
**C.** Cell proliferation assay measured using the CellTiter-Glo kit. One-way ANOVA, *$p < 0.05$.
**D.** Cell migration assay performed using a scratch assay. Images were acquired every 3 hours using MuviCyte, and percentage area coverage was quantified with MuviCyte software. One-way ANOVA, *$p < 0.01$.
**E.** Tumor growth following injection of control and ASF1B knockout SKOV3ip1 cells into nude mice. 1 × 10^6 cells were injected into both flanks of 10 mice for each cell line. Tumor volume was measured twice a week throughout the experiment. One-way ANOVA, **$p < 0.005$.
**F.** Images of subcutaneous tumors collected from the control and knockout groups.

**Suppl. Fig. 5. SYNE1 is downregulated in STIC lesions.**

**A.** H&E (left) and SYNE1 Visium HD spatial expression overlay (right).

**B.** SYNE1 immunohistochemistry. The bottom panels show a higher-magnification view of the boxed regions in the upper panels. In normal fallopian tube cells, SYNE1 localizes to the nuclear envelope, whereas this localization is disrupted in STIC cells.

**Suppl. Fig. 6. MUC6 is increased in postmenopausal fallopian tube epithelium.**

**A.** Uniform Manifold Approximation and Projection (UMAP) of single-cell RNA-sequencing analysis of epithelial cells from 8 premenopausal and 9 postmenopausal nonmalignant fallopian tubes.

**B-D.** Heatmap and boxplot of log2 expression values for SCGB1A1 (B), MUC6 (C), and MUC16 (D). Statistical significance was assessed by ANOVA; p-values are indicated.

**E.** Boxplots of MUC6 and MUC16 expression in bulk benign fimbrial samples from 24 premenopausal and 48 postmenopausal women (GSE297643). Box plots indicate lower quartile, median, upper quartile; whiskers, minimum, maximum. Statistical significance was calculated using the Mann-Whitney U test. p-values are indicated.

**Suppl. Fig. 7. Collagen fiber density is reduced in the stroma adjacent to STIC lesions.**

**A.** Genes overrepresented in STIC-associated compared with FTE-associated stroma.

**B.** Representative Masson's trichrome–stained section highlighting stromal regions adjacent to STIC and FTE.

**C.** Representative ROIs of FTE- and STIC-adjacent stroma with collagen fibers outlined in yellow.

**D.** Computational analysis of collagen-associated H&E image features in the discovery set. After false dicovery correction using the BH method at 0.05, fiber density (FibDens) remained statistically significant.

**E.** Spaghetti plot showing within-case changes in fiber density between matched FTE-adjacent and STIC-adjacent stromal ROIs.

**F.** Forest plot of paired effect sizes with bootstrapped confidence intervals across cases for FibDens feature.

**G.** Median FibDens values measured in digitized H&E slides across stromal ROIs adjacent to HGSC, STIC, and FTE. Differences were assessed using the Mann-Whitney U test.

**Suppl. Fig. 8. Spatial niche and ligand-receptor analyses.**

**A.** Spatial scatter of cells by their niche identity.

**B.** Broad niches defined by cell type density, with niche enrichment for STIC versus FTE. One niche was STIC-enriched and two niches were FTE-enriched relative to global expected levels.

**C.** Cell count of target cells (T cells, macrophages, fibroblasts) proximal to FTE and STIC cells (upper triangles) and target cell-proximal FTE/STIC cells (lower triangles).

### Suppl. Fig. 1

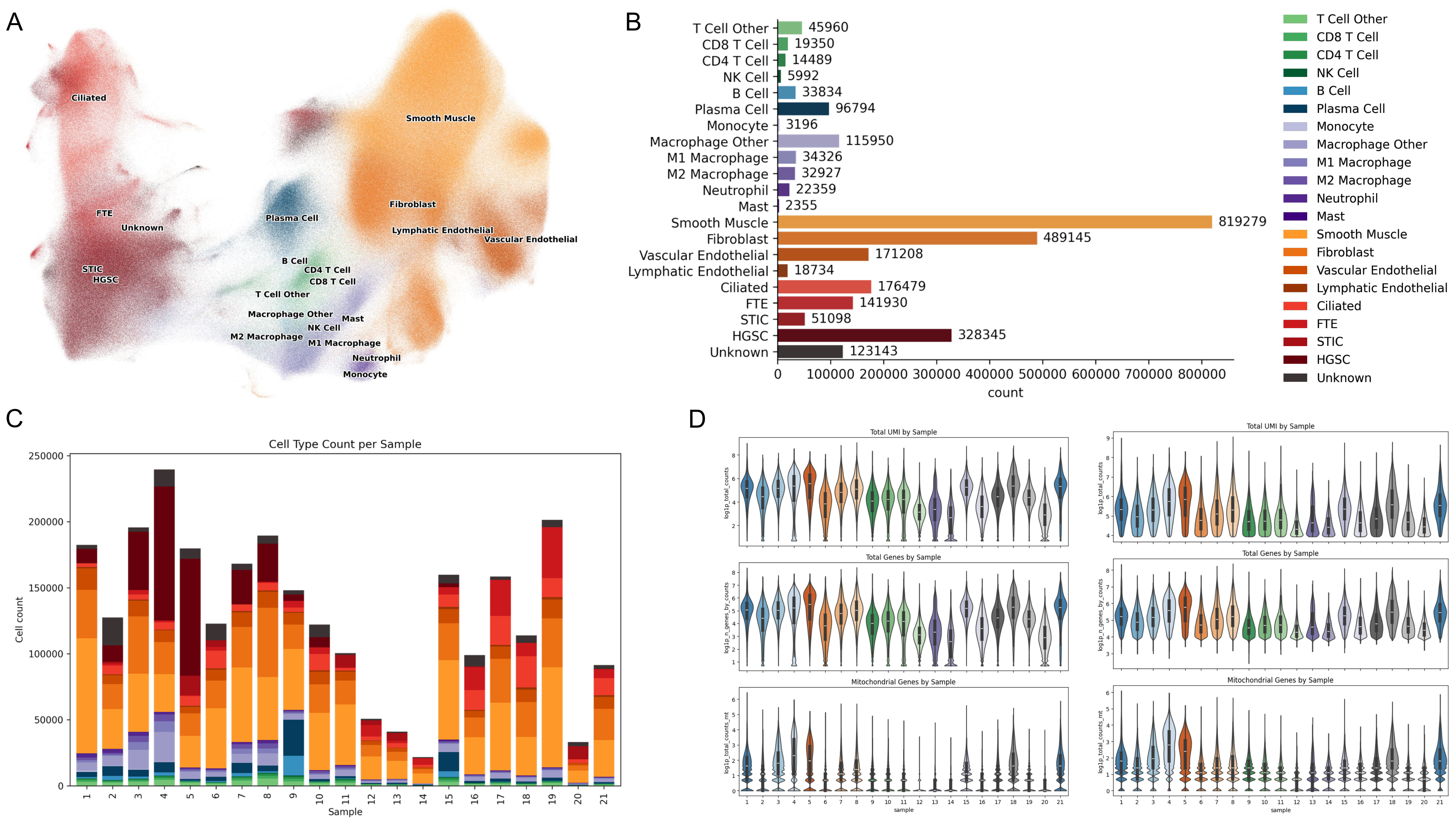
A
Ciliated
Smooth Muscle
FTE
Unknown
Plasma Cell
Fibroblast
Lymphatic Endothelial
Vascular Endothelial
STIC
HGSC
B Cell
CD4 T Cell
CD8 T Cell
T Cell Other
Macrophage Other
Mast
NK Cell
M2 Macrophage
M1 Macrophage
Neutrophil
Monocyte
B
T Cell Other 45960
CD8 T Cell 19350
CD4 T Cell 14489
NK Cell 5992
B Cell 33834
Plasma Cell 96794
Monocyte 3196
Macrophage Other 115950
M1 Macrophage 34326
M2 Macrophage 32927
Neutrophil 22359
Mast 2355
Smooth Muscle 819279
Fibroblast 489145
Vascular Endothelial 171208
Lymphatic Endothelial 18734
Ciliated 176479
FTE 141930
STIC 51098
HGSC 328345
Unknown 123143
0 100000 200000 300000 400000 500000 600000 700000 800000
count
C
Cell Type Count per Sample
Cell count
Sample
D
Total UMI by Sample
Total Genes by Sample
Mitochondrial Genes by Sample
log1p_total_counts
log1p_n_genes_by_counts
log1p_total_counts_mt
sample

#### Suppl. Fig. 2

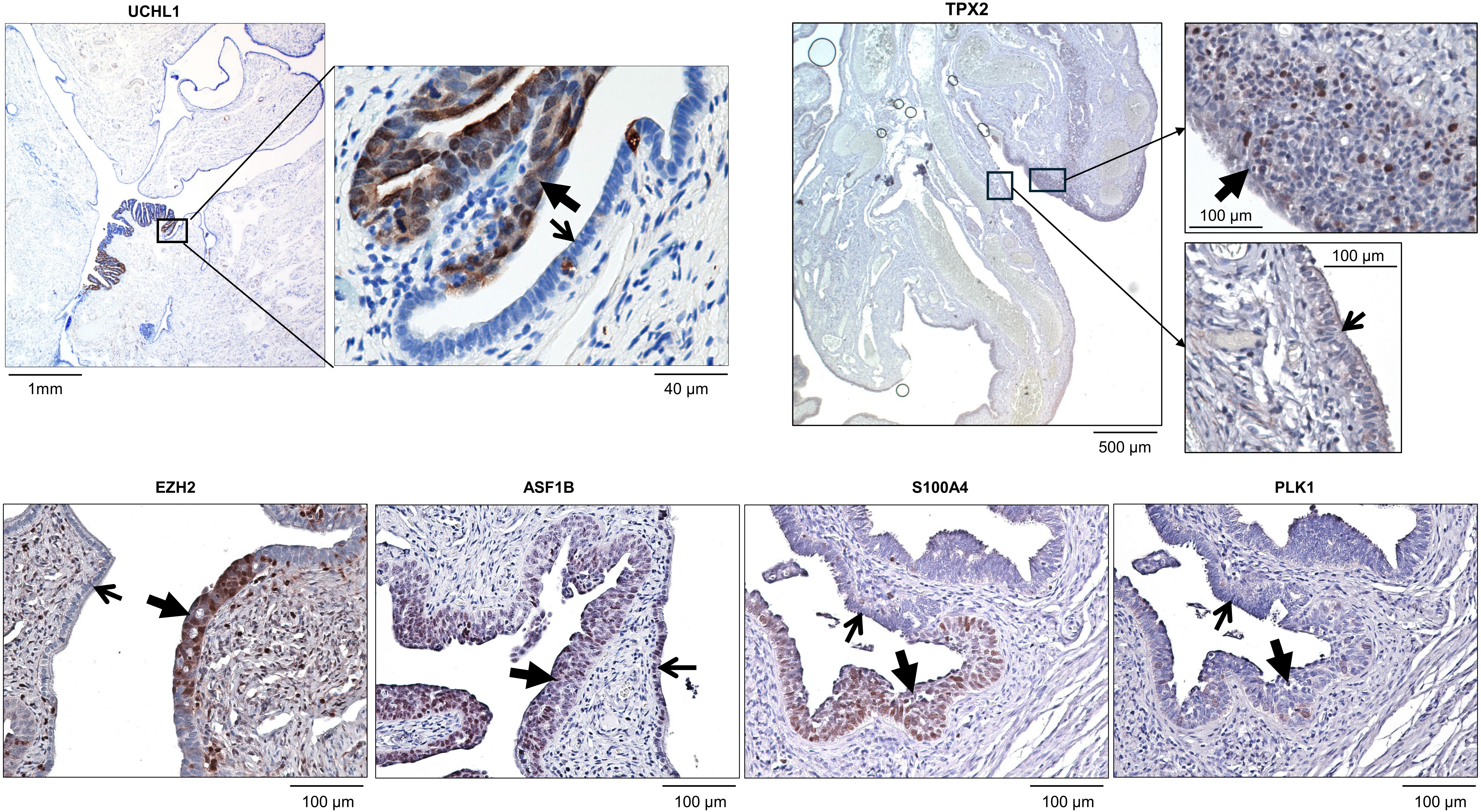
UCHL1
1mm
40 µm
TPX2
100 µm
100 µm
500 µm
EZH2
100 µm
ASF1B
100 µm
S100A4
100 µm
PLK1
100 µm

**Suppl. Fig. 3**

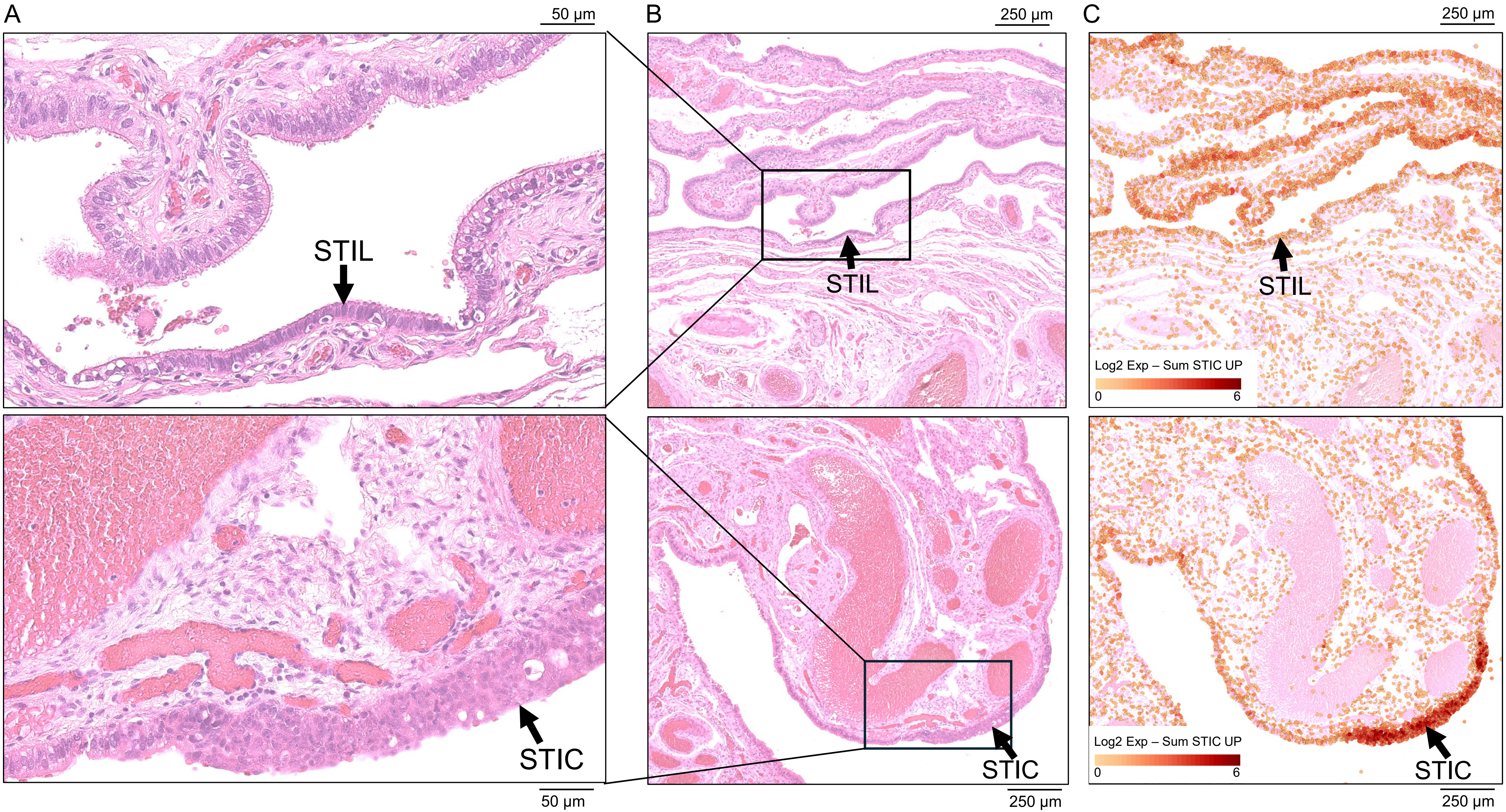
A
50 µm
STIL
STIC
50 µm
B
250 µm
STIL
STIC
250 µm
C
250 µm
STIL
Log2 Exp – Sum STIC UP
0
6
Log2 Exp – Sum STIC UP
0
6
STIC
250 µm

**Suppl. Fig. 4**

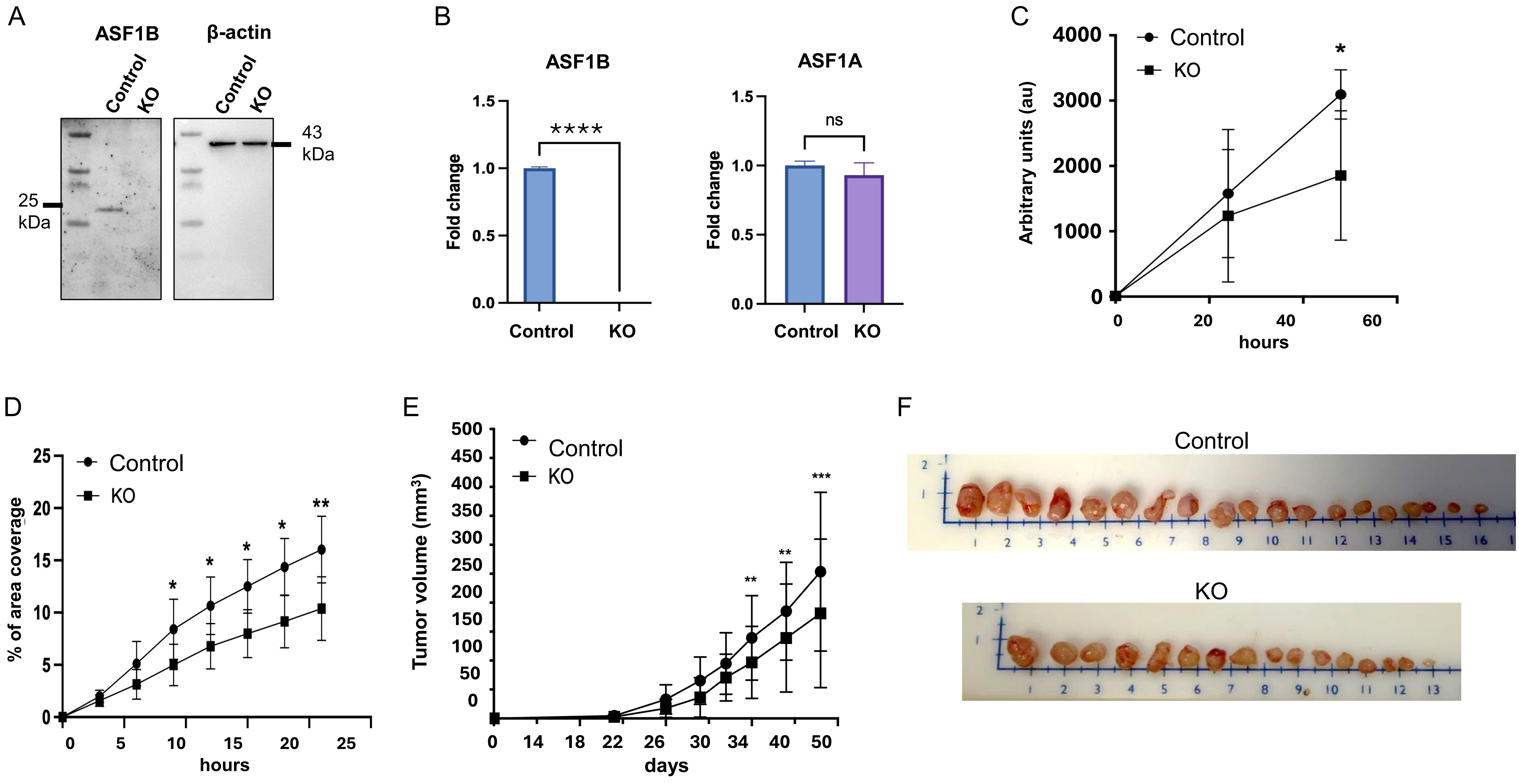
A
ASF1B
β-actin
Control
KO
Control
KO
43 kDa
25 kDa
B
ASF1B
ASF1A
Fold change
1.5
1.0
0.5
0.0
****
ns
Control
KO
C
Control
KO
*
Arbitrary units (au)
4000
3000
2000
1000
0
0
20
40
60
hours
D
Control
KO
% of area coverage
25
20
15
10
5
0
*
*
*
*
**
0
5
10
15
20
25
hours
E
Control
KO
Tumor volume (mm³)
500
450
400
350
300
250
200
150
100
50
0
**
**
***
0
14
18
22
26
30
34
40
50
days
F
Control
KO

**Suppl. Fig. 5**

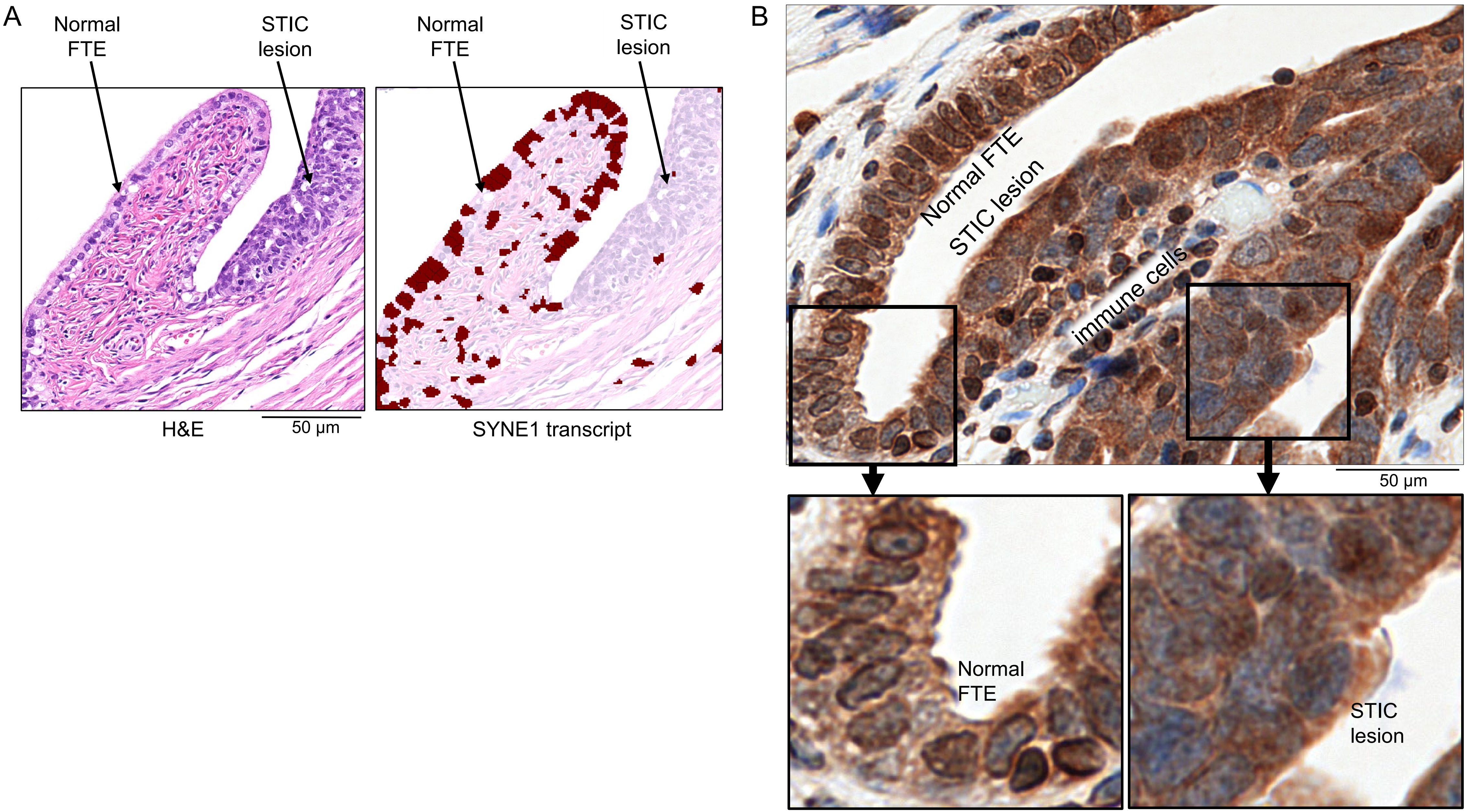
A
Normal
FTE
STIC
lesion
Normal
FTE
STIC
lesion
H&E
50 μm
SYNE1 transcript
B
Normal FTE
STIC lesion
immune cells
50 μm
Normal
FTE
STIC
lesion

#### Suppl. Fig. 6

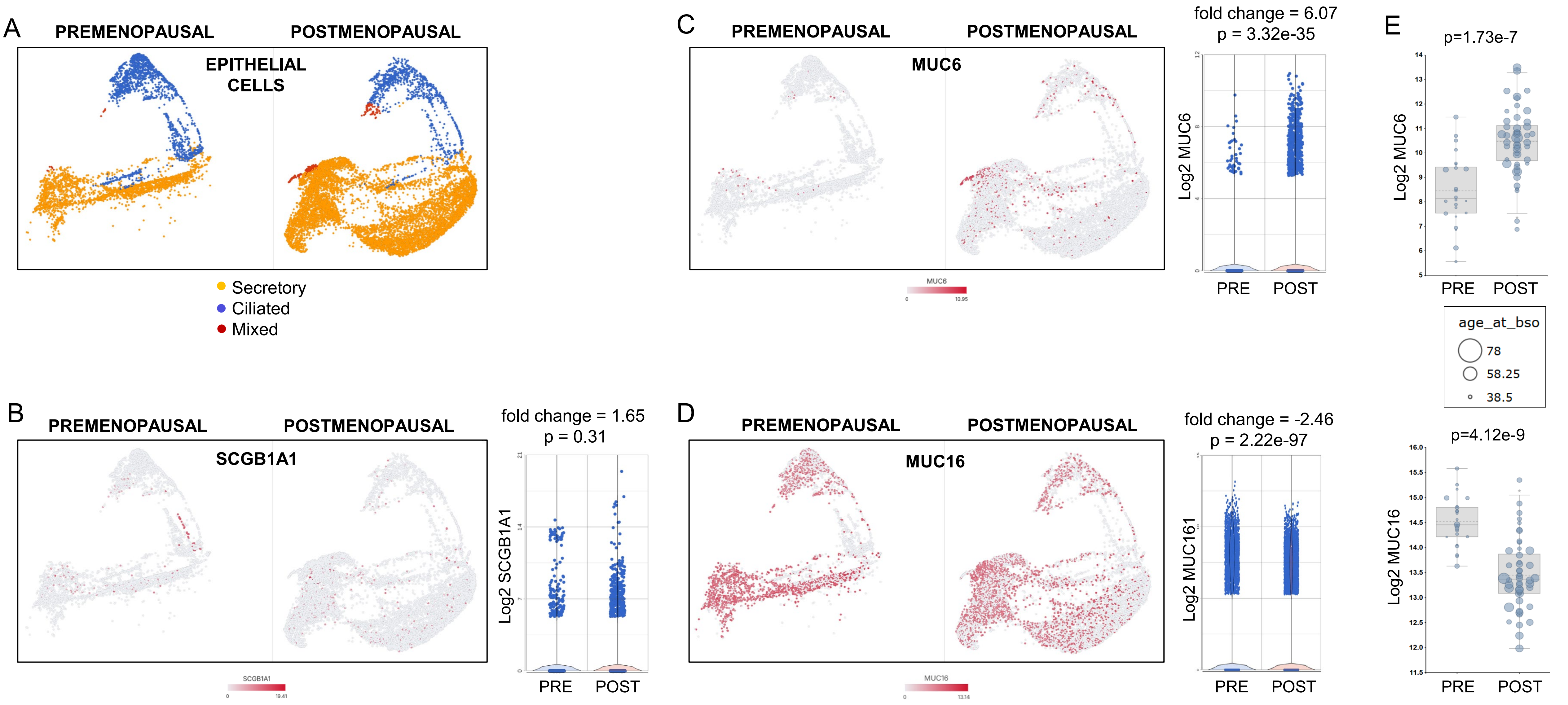
A
PREMENOPAUSAL
POSTMENOPAUSAL
EPITHELIAL CELLS
Secretory
Ciliated
Mixed
B
PREMENOPAUSAL
POSTMENOPAUSAL
SCGB1A1
fold change = 1.65
p = 0.31
Log2 SCGB1A1
PRE
POST
C
PREMENOPAUSAL
POSTMENOPAUSAL
MUC6
fold change = 6.07
p = 3.32e-35
Log2 MUC6
PRE
POST
D
PREMENOPAUSAL
POSTMENOPAUSAL
MUC16
fold change = -2.46
p = 2.22e-97
Log2 MUC161
PRE
POST
E
p=1.73e-7
Log2 MUC6
PRE
POST
age_at_bso
78
58.25
38.5
p=4.12e-9
Log2 MUC16
PRE
POST

### Suppl. Fig. 7

A

| gene | baseMean | log2FoldChange | lfcSE | stat | p-value | p-adj |
|---|---|---|---|---|---|---|
| MGP | 292.7078 | 1.073478 | 0.230637 | 4.654407 | 3.25E-06 | 0.028987 |
| VCAN | 104.8954 | 1.820369 | 0.398406 | 4.569128 | 4.90E-06 | 0.029129 |
| BGN | 69.82845 | 1.404528 | 0.320613 | 4.380755 | 1.18E-05 | 0.052757 |

B

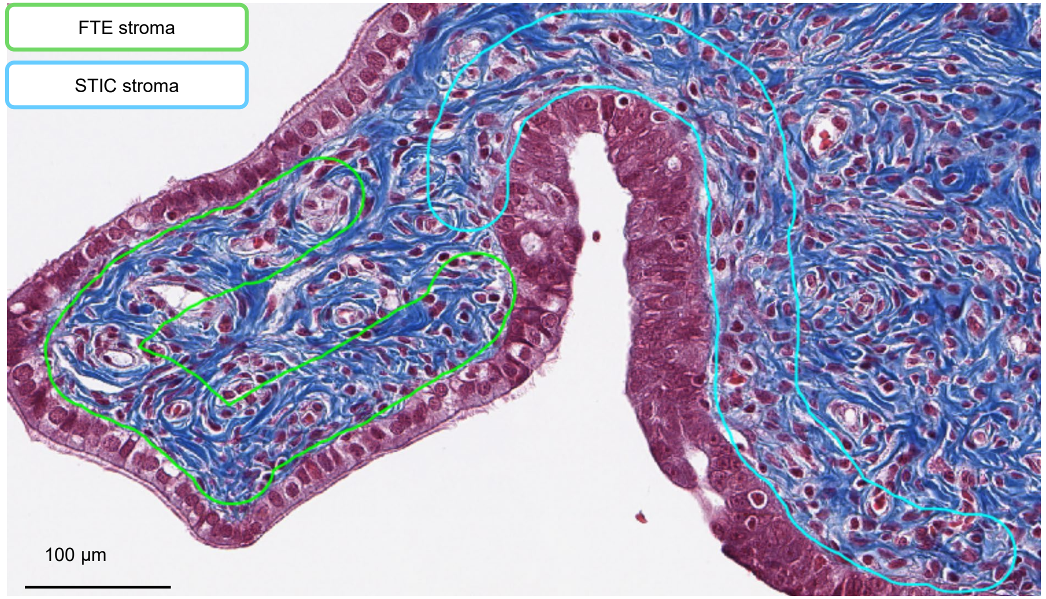


C

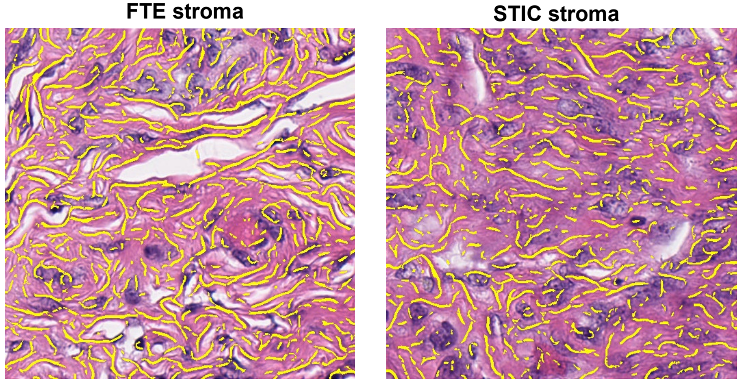


D

| Feature | p-value (raw) | p-value (FDR adj.) | median (STIC* – FTE*) | paired Cohen's d |
|---|---|---|---|---|
| 'FibOrientEntr' | 0.652344 | 0.652344 | -0.01344 | -0.23579 |
| **'FibDens'** | **0.003906** | **0.015625** | **-0.01574** | **-1.36159** |
| 'FibMedLen' | 0.109375 | 0.1484 | -2.75 | -0.70169 |
| 'FibStdLen' | 0.074219 | 0.1458 | -11.6887 | -0.71264 |

* the median feature value across ROIs within a case

E

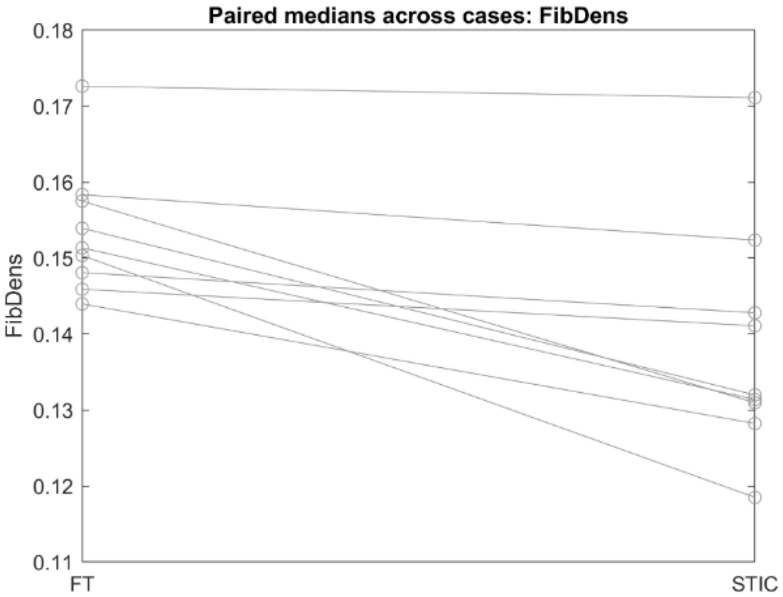


F

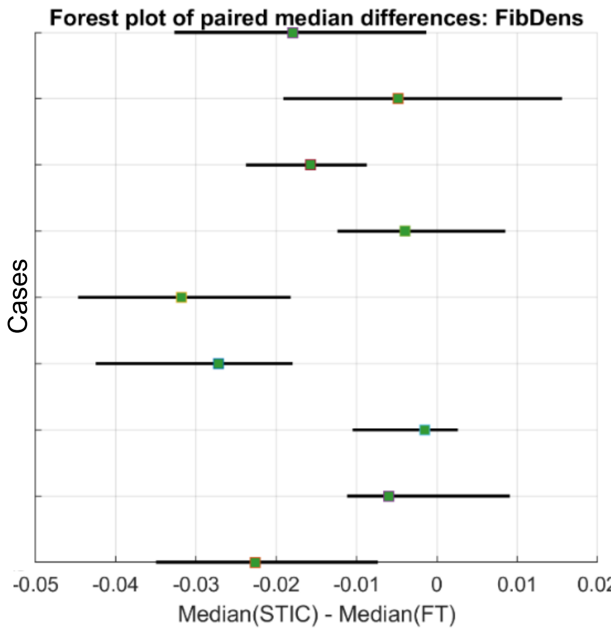


G

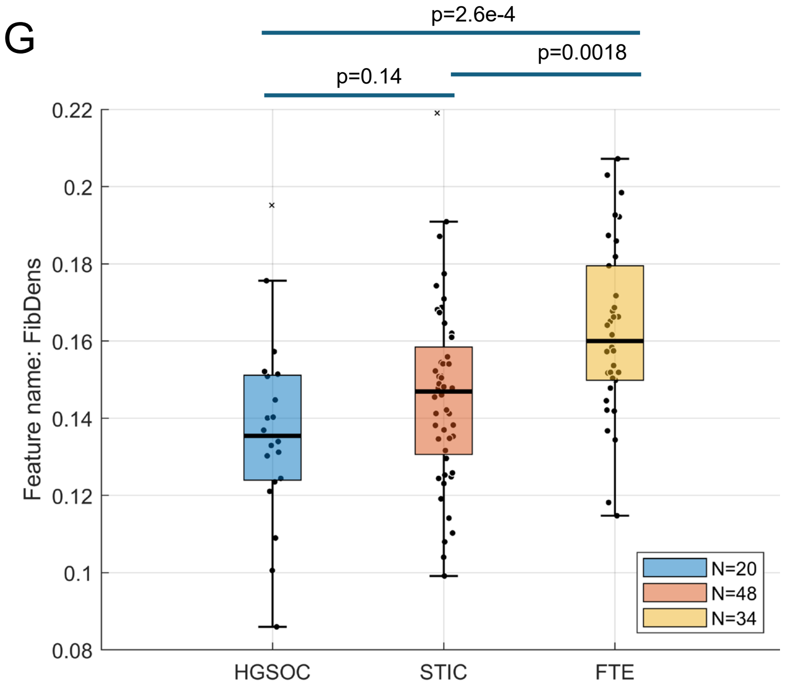

### Suppl. Fig. 8

A

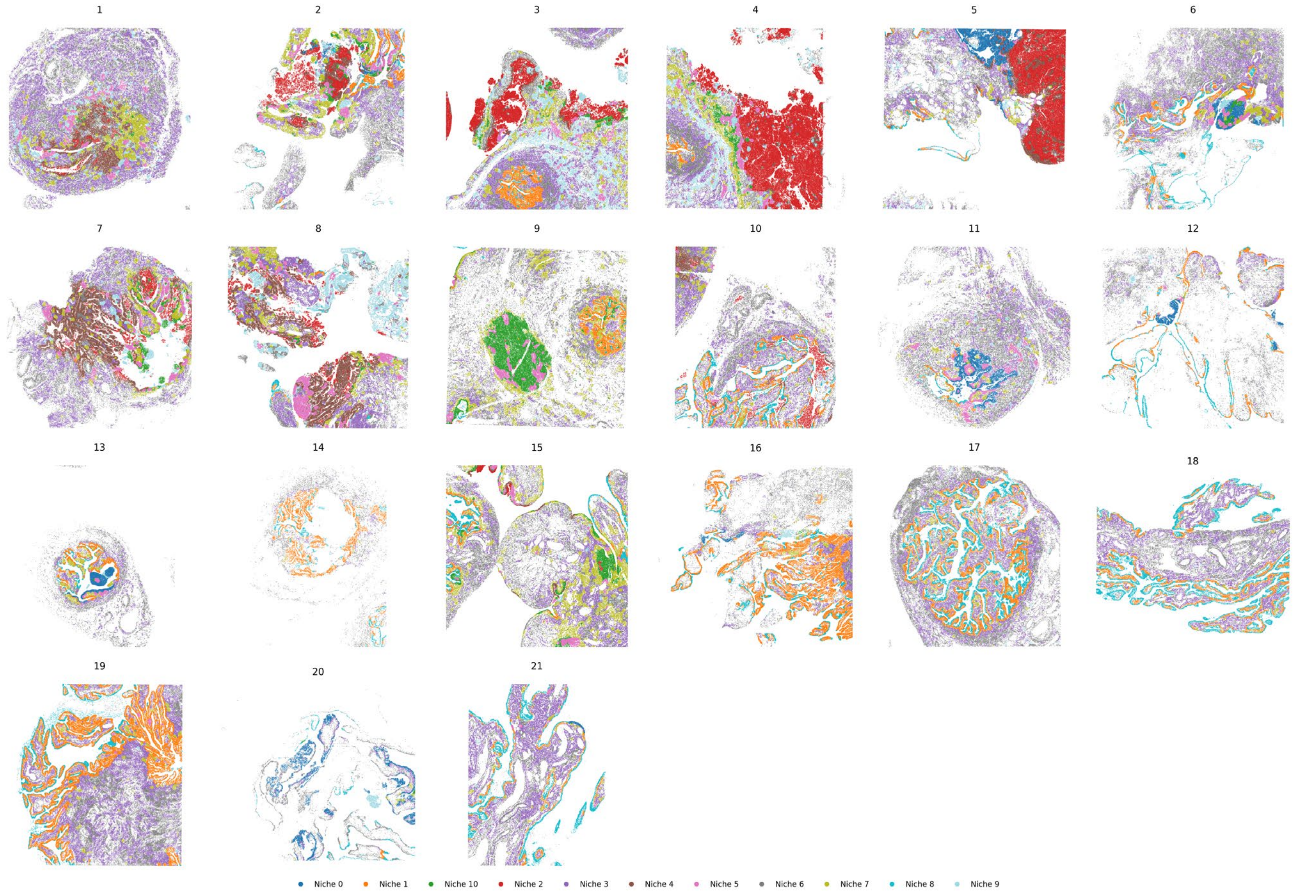


B

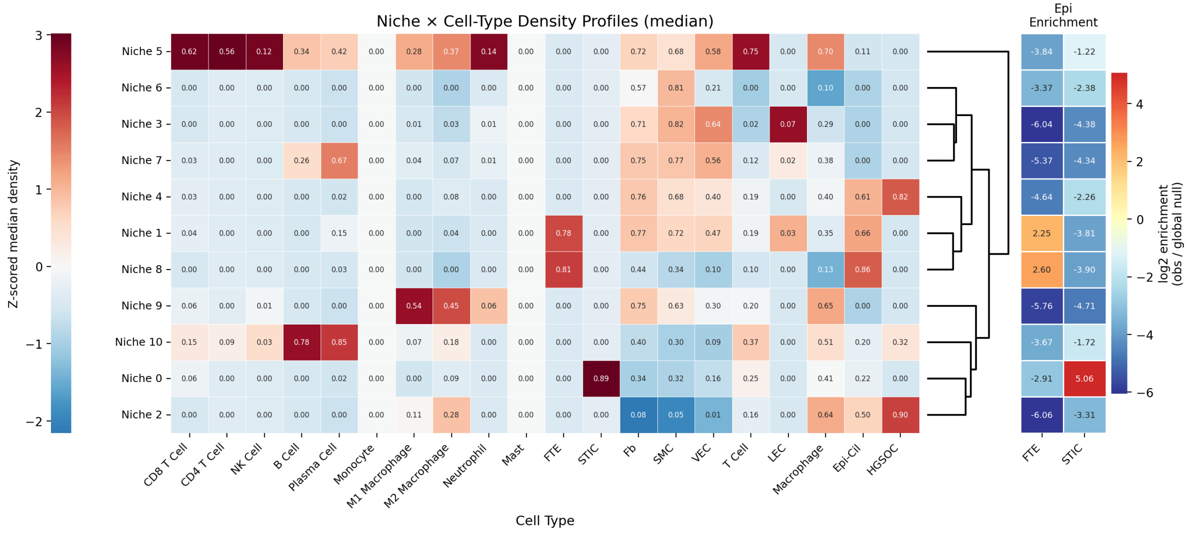


C

| Ligand-receptor analysis cell counts | FTE | STIC |
|---|---|---|
| T cell | 21216 / 3950 | 35480 / 5885 |
| Macrophage | 25423 / 5888 | 38690 / 9798 |
| Fibroblast | 35740 / 29516 | 35257 / 10981 |
